# The Tripod neuron: a minimal structural reduction of the dendritic tree

**DOI:** 10.1101/2022.09.05.506197

**Authors:** Alessio Quaresima, Hartmut Fitz, Renato Duarte, Dick van den Broek, Peter Hagoort, Karl Magnus Petersson

**Affiliations:** Neurobiology of Language Department, Max Planck Institute for Psycholinguistics, 6565 XD Nijmegen, the Netherlands; Donders Institute for Brain, Cognition and Behaviour, 6500 HE Nijmegen, the Netherlands; Faculty of Medicine and Biomedical Sciences, University of Algarve, 8000-810 Faro, Portugal

**Keywords:** dendritic computation, multi-compartmental neuron model, NMDA spikes, spatio-temporal integration, temporal binding

## Abstract

Neuron models with explicit dendritic dynamics have shed light on mechanisms for coincidence detection, pathway selection, and temporal filtering. However, it is still unclear which morphological and physiological features are required to capture these phenomena. In this work, we introduce the Tripod neuron model and propose a minimal structural reduction of the dendritic tree that is able to reproduce these dendritic computations. The Tripod is a three-compartment model consisting of two segregated passive dendrites and a somatic compartment modeled as an adaptive, exponential integrate-and-fire neuron. It incorporates dendritic geometry, membrane physiology, and receptor dynamics as measured in human pyramidal cells. We characterize the response of the Tripod to glutamatergic and GABAergic inputs and identify parameters that support supra-linear integration, coincidence-detection, and pathway-specific gating through shunting inhibition. Following NMDA spikes, the Tripod neuron generates plateau potentials whose duration depends on the dendritic length and the strength of synaptic input. When fitted with distal compartments, the Tripod neuron encodes previous activity into a dendritic depolarized state. This dendritic memory allows the neuron to perform temporal binding and we show that the neuron solves transition and sequence detection tasks on which a single-compartment model fails. Thus, the Tripod neuron can account for dendritic computations previously explained only with more detailed neuron models or neural networks. Due to its simplicity, the Tripod model can be used efficiently in simulations of larger cortical circuits.

## Introduction

Biological neurons integrate complex afferent inputs within a dendritic structure which accounts for most of the spatial extent of a neuron. The dendritic arborization hosts a significant part of excitatory and inhibitory synapses and processes the input signals before the resulting signal reaches the cell body, and in particular the axon hillock. In the dynamical systems theory of neural information processing, neurons function as non-linear, non-stationary (and stochastic) operators, and the dendrites determine important aspects of the neurons’ transfer characteristics (Payeur *et al*., 2021; Stuart & Spruston, 2015; Gidon *et al*., 2020; Poirazi & Papoutsi, 2020; Larkum *et al*., 2022).

Neuron models that explicitly consider the dynamics of the dendritic tree are typically referred to as multi-compartment models. These models capture the spatio-temporal dendritic dynamics by introducing additional state variables and differential equations which describe the dynamics of the dendritic membrane potential (Koch, 1999). Depending on the implemented dendritic architecture, membrane dynamics, and receptors/ion-channel repertoire, high-resolution multi-compartmental models can reproduce the membrane physiology in detail (Winnubst & Lohmann, 2012; Branco *et al*., 2010; Ujfalussy *et al*., 2018). Simulations with neuron models including dendrites shed light on important problems of brain functions, including unsupervised learning (Bono & Clopath, 2017; Payeur *et al*., 2021), signal filtering (Yang *et al*., 2016), temporal discrimination (Branco *et al*., 2010), coincidence detection (Mel, 1992; Poirazi *et al*., 2003), structured sequence processing (Haga & Fukai, 2018; Ahmad & Hawkins, 2016), and the creation and maintenance of associative memories (Kastellakis *et al*., 2016). This body of evidence suggests that dendritic processing is fundamental to nervous system computation. However, the computational cost of simulating detailed multi-compartment models impedes their use in large networks. Thus, most studies that analyze processing properties in large networks do not explicitly consider dendritic structure but often use simpler point neuron models instead. These studies regard neural computation as the outcome of the particular network structure used, disregarding the complexity of cell-internal processes (Duarte & Morrison, 2019; Potjans & Diesmann, 2014; Haeusler *et al*., 2009; Bastos *et al*., 2012).

The present work introduces a computationally efficient, three-compartment model that includes relevant dendritic degrees of freedom and remains simple enough to be used in larger network simulations. This model, which we call the Tripod neuron, is derived from previous theoretical and experimental work, and three main ingredients define its dynamics. First, the Tripod has two dendritic compartments. This is the minimum number of dendritic compartments, in addition to the somatic compartment, which allows a branching dendritic tree. Several studies have shown that relatively few dendritic degrees of freedom are sufficient to reproduce the nonlinear integration effects of apical dendrites in pyramidal cells (Poirazi *et al*., 2003; Larkum, 2013). Accordingly, an extensive comparison of the number of dendritic compartments to mimic *in-vivo* dynamics indicates that two compartments are sufficient to explain most of the observed variability in the somatic membrane potential (Ujfalussy *et al*., 2018; Wybo *et al*., 2021), and models with more than two dendritic compartments show modest qualitative differences (Ahmad & Hawkins, 2016; Bono & Clopath, 2017; Kastellakis *et al*., 2016). Secondly, the internal dynamics of the Tripod neuron is consistent with observed neurophysiology. The dendritic structure consists of two isolated compartments connected to the somatic compartment. Each compartment integrates fast and slow excitatory and inhibitory inputs locally through conductance-based synapses, and we show that a simple circuit approximation (Koch, 1999) suggests that a single degree of freedom, the electrotonic distance from the soma, determines an integration time-scale of the dendrites and analytically defines two types of compartments, here called short and long dendrites. Finally, we investigated slow voltage-dependent NMDA receptors that mimic an important property of dendritic computation. When the post-synaptic potential exceeds a certain threshold, the NMDA receptors open to Ca^2+^ ions and boost post-synaptic membrane depolarization, generating a so-called NMDA spike, or plateau potential (Antic *et al*., 2010; Mel, 1992; Tabone & Ramaswami, 2012). This non-linear phenomenon, along with self-regenerative events such as back-propagating spikes (Rapp *et al*., 1996) in proximal dendrites, enrich the computational toolkit of the dendrites and determine the most interesting properties of the present model. The slow voltage decay of the dendritic potential provides a short-term dendritic memory which is not accounted for by other adaptation mechanisms in single-compartment models, for example (Fitz *et al*., 2020; Brette & Gerstner, 2005). This aspect of our work complements previous studies of NMDARs in models with a small number of compartments (Bono & Clopath, 2017; Mel, 1992; Yang *et al*., 2016), and provides a basis for further explorations of the role of NMDA spikes in neuronal working memory (Fitz *et al*., 2020; Wang, 1999), and temporal binding (Augusto & Gambino, 2019; Baggio & Hagoort, 2011).

## Methods

### The Tripod neuron model

The *Tripod* neuron is composed of three separate computational elements, or compartments. It has an axosomatic compartment, representing the soma and perisomatic locations, and two electrotonically segregated dendritic compartments coupled to the soma in a Y-shape (Fig. 1).

**Figure 1:**
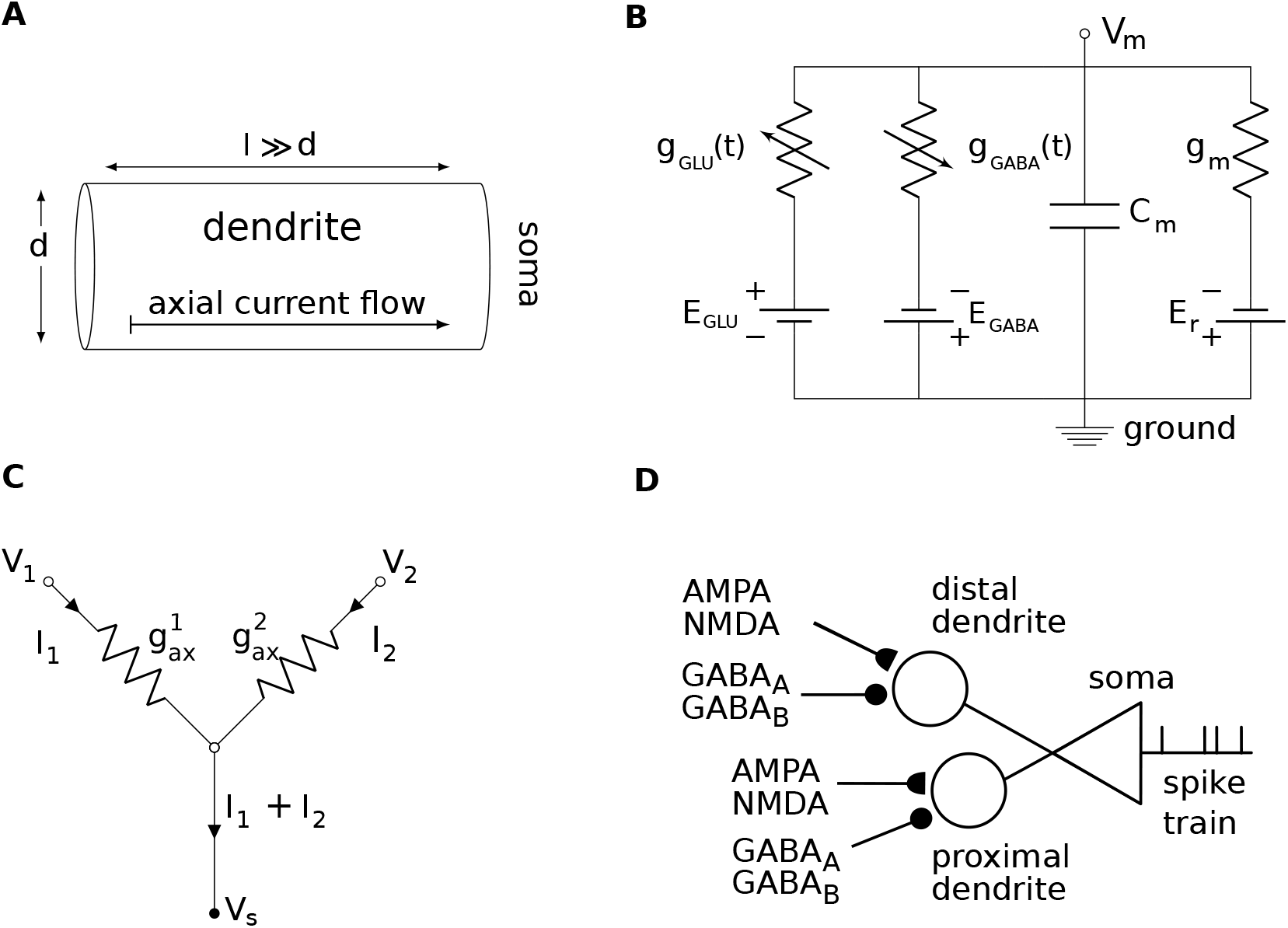
Schematic of the Tripod neuron. **(A)** Dendritic compartments were modeled as a cylindrical segment of a cable with length *l* and diameter *d*. Their electrical properties were set by the membrane patch equations (Eqs. 5, 6, 7) and membrane-specific parameters (Table 2). When dendrites had a larger potential than the soma, current flowed along the dendritic axis towards the soma. **(B)** Circuit diagram of a dendritic membrane patch with time-varying conductances across the membrane. Conductances were regulated by glutamatergic receptors *g*_GluRs_ or GABAergic receptors *g*_GABA_ with reversal potentials *E*_GluRs_ and *E*_GABA_, respectively (Table 4). The membrane reversal potential *E*_*r*_ coupled in series with the leak conductance *g*_*m*_ and the membrane acted as a capacitance *C*_*m*_ with respect to the extracellular space (ground). The membrane potential *V*_*m*_ was determined by the currents flowing to the dendritic compartment. **(C)** The dendritic potentials *V*_1_ and *V*_2_ were coupled to the somatic membrane *V*_*s*_ through the axial conductances 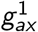 and 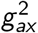. The resulting current *I*_1_ + *I*_2_ flowed dromically from the dendrites to the soma. **(D)** The Tripod neuron with two dendrites and a somatic compartment. Each dendrite received synaptic input mediated by four types of receptors, AMPA, NMDA, GABA_A_ and GABA_B_. Distal dendritic compartments were modeled using a smaller axial conductance compared to proximal ones. The spike-generating soma is represented as a triangle.

#### Axosomatic compartment

The soma was modeled as an adaptive exponential integrate-and-fire (AdEx) neuron (Brette & Gerstner, 2005). It is a two-dimensional neuron that models the dynamics of the somatic membrane potential *V* ^*s*^ and an adaptive current *w* :

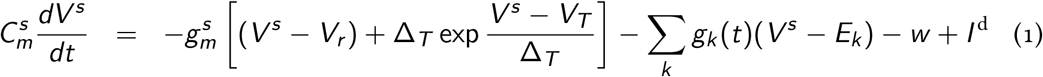

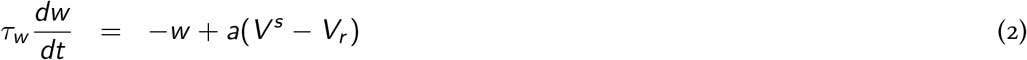

The leak conductance 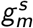 defines the permeability of the somatic membrane, 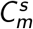 its capacitance and *g*_*k*_ the set of variable synaptic conductances (Fig. 1*B*). The synaptic conductances and reversal potentials *E*_*k*_ are further described in the section *Synaptic dynamics* below. We use the superscript *s* throughout to refer to variables and parameters of the somatic compartment, whereas the superscript *d* refers to dendritic compartments. The first equation of the AdEx neuron aims to reproduce the sub-threshold and spike-onset dynamics of pyramidal cells. For a membrane potential *V* ^*s*^ below the rheobase threshold *V*_*T*_, the neuron behaves as a leaky integrator of the currents from the dendritic compartments *I*_*d*_ and the somatic leakage conductances *g*_*k*_ (*V* ^*s*^ − *E*_*k*_). For larger depolarizing events, the membrane potential exceeds the rheobase threshold *V* ^*s*^ *> V*_*T*_ and activates the exponential non-linearity, mimicking a spike-generation mechanism. The slope of the exponential growth is governed by Δ_*T*_. The spike events occur at times *t*^*f*^ when *V* ^*s*^ exceeds the spiking threshold *u*_*th*_. Afterwards the membrane potential is reset to *V*_*r*_ and the adaptation current *w* is increased by a constant value *b*. The adaptation current accounts for several physiological processes and decreases the excitability of the neuron after it has spiked. All parameters of the somatic compartment were fixed and set to the values used in Brette & Gerstner (2005), except for the somatic leak conductance which was set to 40 nS in agreement with the multi-compartment model of Bono & Clopath (2017), see Table 1. The reset potential of the AdEx model has been set to *u*_*r*_ = −70.6 mV as in (Brette & Gerstner, 2005) rather than to *u*_*r*_ = −55 mV (Duarte & Morrison, 2019; Bono & Clopath, 2017) so that the bursting behavior in the Tripod will depended only on the dendritic dynamics.

**Table 1:** Parameters for the axosomatic compartment of the Tripod neuron. Values corresponds to those proposed in Brette & Gerstner (2005), except for the somatic leak conductance which is set to 40 nS, as in Bono & Clopath (2017).

| Symbol | Description | Value | Unit |
| --- | --- | --- | --- |
| $g_L$ | Membrane leak conductance | 40 | nS |
| $C_m$ | Membrane capacitance | 281 | pF |
| $V_r$ | Resting membrane potential | -70.6 | mV |
| $V_T$ | Threshold potential | -50.4 | mV |
| $u_{th}$ | Spike onset threshold | 0 | mV |
| $u_r$ | Reset potential | -70.6 | mV |
| $\Delta_T$ | Slope factor | 2 | mV |
| $\tau_w$ | Spike-triggered adaptation time scale | 144 | ms |
| $a$ | Subthreshold adaptation conductance | 4 | nS |
| $b$ | Spike-triggered adaptation increment | 80.5 | pA |
| $t_{up}$ | Spike width (soma clamped at 20 mV) | 1 | ms |
| $t_{ref}$ | Refractory period | 2 | ms |

**Table 2:** Dendritic physiology parameterized for human and mouse, following Koch (1999) and Eyal *et al*. (2016).

| Symbol | Description | Human | Mouse | Unit |
| --- | --- | --- | --- | --- |
| $r_m$ | Membrane resistance | 39 | 1.7 | $\text{k}\Omega \text{ cm}^2$ |
| $r_{ax}$ | Intracellular resistance | 200 | 200 | $\Omega \text{ cm}$ |
| $c_m$ | Membrane capacitance | 0.5 | 1 | $\mu\text{F}/\text{cm}^2$ |
| $V_r$ | Resting potential | -70.6 | -70.6 | mV |

#### Dendritic compartments

Dendritic compartments were approximated as conductive cylinders whose voltage was governed by a passive membrane-patch equation similar to the soma but lacking mechanisms for spike generation and intrinsic adaptation:

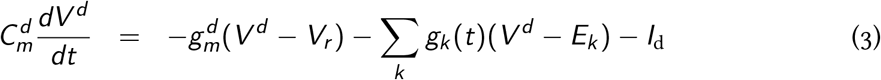

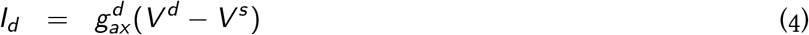

The current *I*_*d*_ was computed as the potential difference between the dendritic and somatic compartment, multiplied by the axial conductance 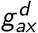 (Fig. 1*C*). Current flow was positive from the dendrites to the soma, *I*_*d*_ *>* 0, except when the somatic potential *V*_*s*_ exceeded the firing threshold and the neuron emitted a spike. Consistent with Bono & Clopath (2017), we captured the backpropagation of somatic action-potentials by clamping *V*_*s*_ (*t*^*f*^) to 20 mV for 1 ms. The effect of the backprogating action potential is illustrated in Fig. 2**D**.

**Figure 2:**
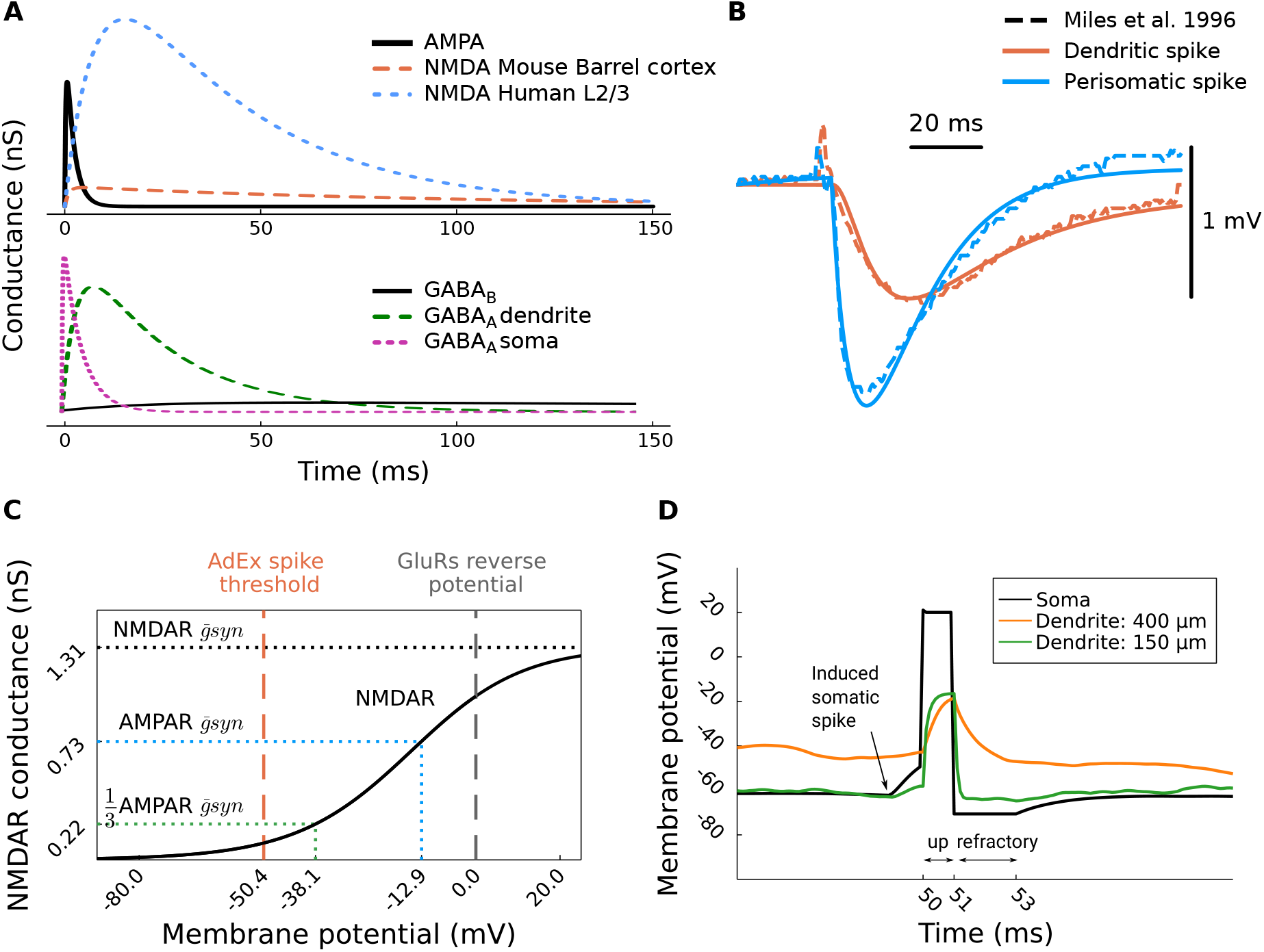
Synaptic kinetics and conductance, and backpropagating action potential. **(A)** The dynamics of glutamatergic (upper panel) and GABAergic (lower panel) synapses for the parameters reported in Table 4. **(B)** Fit of GABA_A_ timescale and maximal conductance for somatic and dendritic synapses, original data (dashed line) from Miles *et al*. (1996). **(C)** NMDAR conductance as function of the compartment membrane potential. Horizontal dotted lines express the voltage-independent conductance of AMPARs and the maximal NMDARs conductance. **(D)** Back-propagating action potential in the dendrites. The backpropagation is purely due to the high membrane potential of the somatic compartment during the spike duration (1 ms). After reset the membrane potential of the soma is held fixed at the reset potential (*u*_*r*_) for the entire duration of the refractory period (2 ms).

#### Dendritic geometry

The capacitance 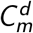, leak conductance 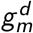, and axial conductance 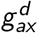 of the dendritic compartments depended both on the geometry and the membrane properties.

The macroscopic parameters 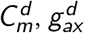, and 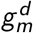 can be computed from the relative densities *c*_*m*_, *r*_*ax*_ and *r*_*m*_ via the standard cable theory (Koch, 1999):

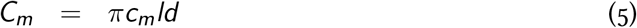

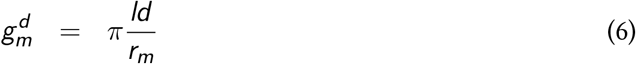

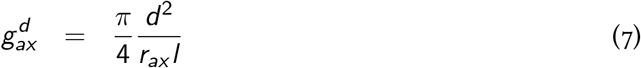

where *l* and *d* refer to the length and diameter of the dendritic cylinder (Fig. 1*A*), respectively. The microscopic parameters *c*_*m*_ and *r*_*m*_ reflect the trans-membrane capacitance and resistance per unit of surface area and *r*_*ax*_ the axial resistance per units of volume that a dendritic current experiences in the direction of the axosomatic compartment. The integration timescale *τ*_*d*_ of a dendritic compartment is given by the effective timescale of the corresponding RC circuit:

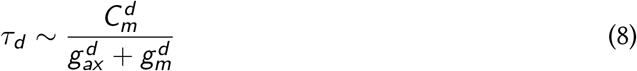

#### Synaptic dynamics

For synaptic transmission we considered the principal receptors concerning excitation and inhibition, including two glutamatergic receptors with fast (AMPA) and slow (NMDA) dynamics, and two GABAergic receptors with short (GABA_A_) and long (GABA_B_) timescales. Each receptor was modeled as a conductance with double-exponential kinetics (Roth & van Rossum, 2009):

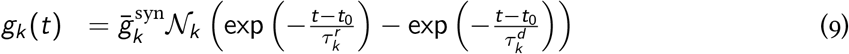

with *k* ∈ {AMPA, NMDA, GABA_A_, GABA_B_} indicating that each receptor has specific parameters. The equation describes the rise and decay of the receptor conductance *g*_*k*_. The timescale of rise and decay are give by *τ*_*r*_ and *τ*_*d*_ while the amplitude of the curve is defined by the maximal conductance parameter *g*_*syn*_. To ensure that the amplitude equals 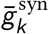, the conductance was scaled by the fixed normalization factor *N*_*k*_. This normalization factor is computed, for each receptor type, as

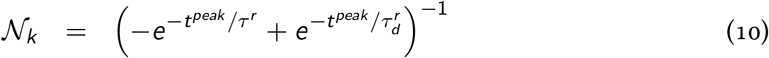

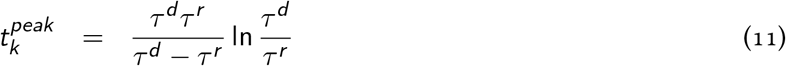

The ratio between the maximal conductance of the NMDA and the AMPA receptor is defined as NMDA-to-AMPA ratio (NAR). The conductance gating of the NMDAR depends on the intracellular depolarization which is captured by a multiplicative voltage-gating mechanism:

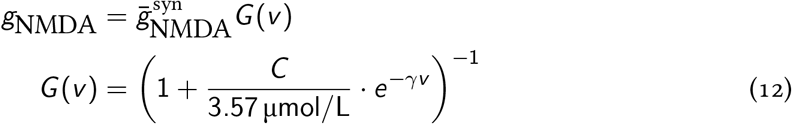

where *γ* regulates the steepness of the voltage-dependence. The extracellular concentration *C* of magnesium ions Mg^2+^ was fixed at 1 μmol*/*L. These equations and parameters were obtained from Jahr & Stevens (1990). The rise and decay timescales of the NMDAR, the NAR, and *γ* assume different values in mouse (Duarte & Morrison, 2019) and human neurophysiology (Eyal *et al*., 2018). All compartments were endowed with excitatory and inhibitory synapses but differed in relative receptor composition and the corresponding parameters. Following previous experimental findings (Petralia *et al*., 1994; Schulz *et al*., 2018) and modelling work (Pongrácz *et al*., 1992), NMDARs were located only on the dendritic compartments. However, this was inconsequential in the Tripod model because the voltage threshold for NMDAR activation was larger than the somatic firing threshold, thus resulting in no contribution of NMDA channels to the somatic synaptic current. During stimulation of glutamatergic receptors, both NMDARs and AMPARs are activated. Even though the NMDAR voltage-dependent component in Eq. 12 is continuous, its non-linear rise allows us to define a soft-threshold at approximately −40 mV. This value is referred to as the NMDA spike-threshold throughout the manuscript. We chose −40 mV because for more hyperpolarized membrane potentials (below the threshold) the NMDAR conductance is less than one third of its AMPAR counterpart and does not trigger NMDA-spikes, as shown in Fig. 2*C*. To parameterize the inhibitory responses, we fit the inhibitory post-synaptic potentials (IPSPs) obtained from guinea-pig hippocampus (Miles *et al*., 1996), which characterize the dendritic versus somatic inhibition on pyramidal cells and can be considered as an effective parametrization of the differences between perisomatic and dendritic inhibition. The timescales obtained from data entail that inhibitory inputs on dendritic compartments have a slower timecourse, whereas somatic inhibitory inputs have a larger amplitude and faster rise and decay, and suggest that somatic GABAergic transmission is mediated primarily by GABA_A_ receptors (Miles *et al*., 1996).

#### Fit of inhibitory synapses

The fit was achieved by reproducing the somatic IPSPs reported in Miles *et al*. (1996). The Tripod neuron was held at resting potential and the inhibitory reversal potential was further lowered of −30 mV, similarly to the experimental procedure used to record the data. The fit was performed on the minimal IPSPs, which correspond to the smallest quanta of PSP that a single inhibitory synapse could elicit in the soma. Considering that the inhibitory neurons stimulated in the physiological experiment had more than a single synaptic contact with the pyramidal cell, we compared the fit to the stimulation of 5 simulated synapses.

### Numerical simulation

Numerical integration used an improved forward Euler method (Heun’s method (Ascher & Petzold, 1998)) with explicit integration and a step-size of 0.1 ms. Dendritic currents were computed from the potential difference between two coupled compartments. Because of the short integration step, the order of integration of dendrites and axosomatic compartments was unimportant. We computed the axial currents first, then the dendritic and somatic voltage changes. Note that the time-step of the explicit integration scheme used is less than half of the fastest timescale in the system, and that the time scales in the model are within two orders of magnitude of each other and the explicit Table 3; therefore, the integration scheme does not incur in numerical instability or stiffness issues at double precision computation that can emerge in the integration of cable equations in fine-grained spatial discretization models (Carnevale & Hines, 2006). Simulations were performed in Julia using custom code which can be obtained on ModelDBLINK and on https://github.com/aquaresima/.

**Table 3:** Parameters for dendritic compartments computed from the physiological specifics in Table 2.

| Symbol | Description | Human |  | Mouse |  | Unit |
| --- | --- | --- | --- | --- | --- | --- |
|  |  | distal | proximal | distal | proximal |  |
| $l$ | Dendritic length | 400 | 150 | 400 | 150 | $\mu\text{m}$ |
| $d$ | Dendritic diameter | 4 | 4 | 4 | 4 | $\mu\text{m}$ |
| $g_m$ | Leak conductance | 1.29 | 0.32 | 29.57 | 7.39 | nS |
| $\tau_d$ | Membrane timescale | 1.48 | 0.22 | 1.11 | 0.36 | ms |
| $g_{ax}$ | Axial conductance | 15.71 | 62.83 | 15.71 | 62.83 | nS |
| $C_m$ | Membrane capacitance | 25.13 | 6.28 | 50.27 | 12.57 | pF |

## Results

Physiological parameters for pyramidal cells are difficult to reconcile across datasets because there exists significant morpho-physiological variation in the mammalian neocortex, both across species and across regions and laminae. The functional consequences of this variation can be difficult to assess. In this section, we show that some of this variability—in particular in the membrane timescale and differences in excitability between human and mouse pyramidal cells—can be explained by explicitly incorporating simple dendritic geometry and membrane physiology. We report important differences in the neuron model behavior when varying the dendritic morphology, the capacitive properties of the cell membrane, and the dendritic NMDA-to-AMPA ratio (NAR).

### Dendritic geometry determines activation boundaries

Excitatory synaptic input to the dendritic tree results in a forward, dromic flow of depolarizing current. This current depends on the potential difference between the perisomatic region and the location of synaptic contact, with an upper bound set by the maximum depolarization that the dendritic compartments can reach. Given the axial resistance and membrane leakage, the geometry of the dendritic branch determines whether dendritic activity can elicit somatic spikes or not. Here, we determine these *activation boundaries* as a function of dendritic length, diameter, and membrane physiology in mouse versus human pyramidal cells. Assuming that a dendrite of the Tripod is fully depolarized after a synaptic event, its capacity to generate somatic spikes is determined by the ratio between the axial conductance 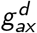 and the membrane leakage 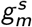 at the soma. For integrate-and-fire neurons, the dendrite can generate a spike when the following equation is satisfied (the full derivation is given in Appendix A):

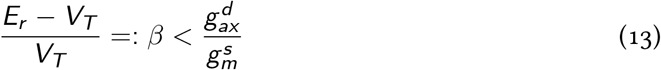

where *E*_*r*_ is the resting membrane potential, *V*_*T*_ is the spike threshold, and 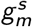 is the leak conductance of the soma. These parameters depend on the somatic compartment. In our model, *β* is constant and the only variable in Eq. 13 is the axial conductance 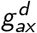 which is determined by Eq. 7 through the cable diameter *d*, its length *l*, and the specific axial resistance *r*_*ax*_ defined by the membrane physiology (Rall, 2011). Hence, Eq. 14 defines a geometrical region where a dendrite can generate a spike. Following similar reasoning, we identify a second geometric region where full depolarization of a single dendrite is insufficient to elicit a somatic spike, but the simultaneous activation of two dendrites can:

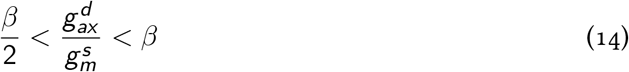

The two regions identified by Eq. 13 and Eq. 14 are shown in blue in Fig. 3*A* and are referred to as *spiking* regions.

**Figure 3:**
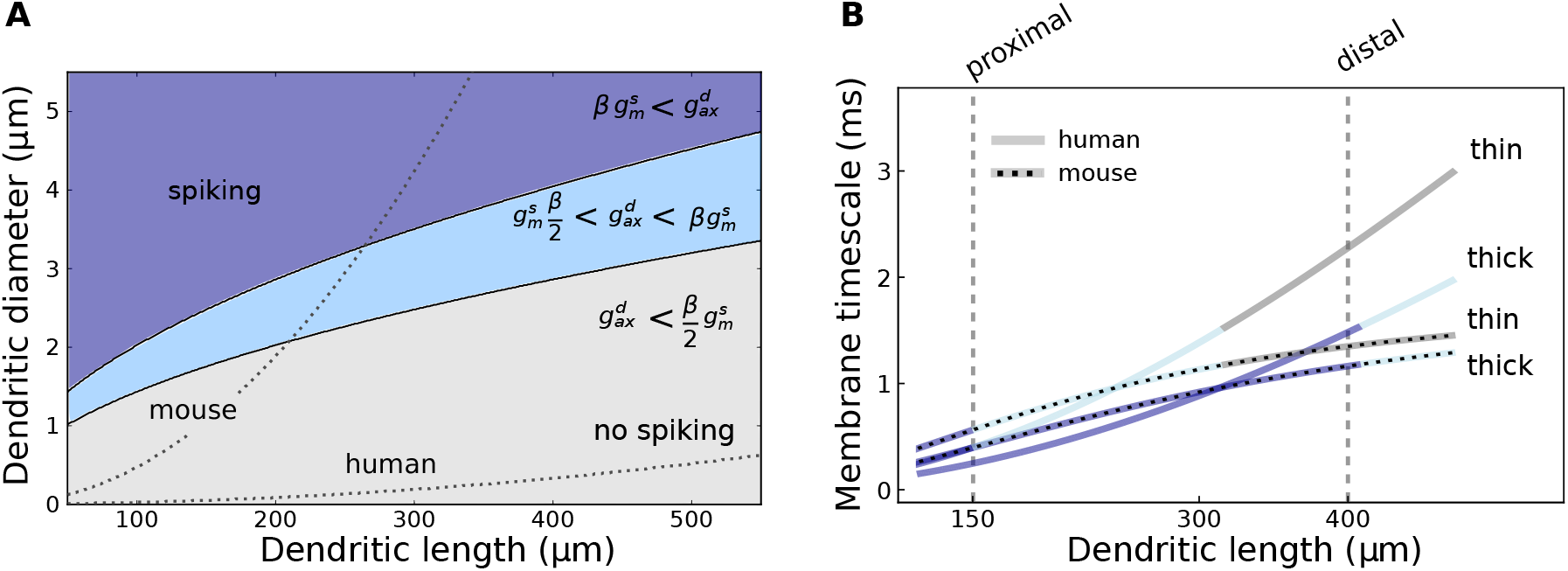
The functional contribution of dendrites to the somatic response depends on dendritic geometry. **(A)** Phase diagram for the axial conductance 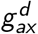 as a function of dendritic diameter and length. Solid black lines show the boundaries imposed by the inequalities of Equation 14. They separate configurations where dendritic depolarization alone cannot elicit somatic spikes (grey region), only co-active dendritic compartments elicit somatic spikes (light blue), or depolarization of a single dendrite can elicit somatic spikes (dark blue). The geometrical regions for spike-onset onset are computed assuming the compartments clamped at *E*_GluRs_, as described in AppendixA; because the specific axial conductance is similar for human and mouse cells, there are no species specific differences in the geometries that lead to somatic spikes. Dotted lines mark the boundaries above which 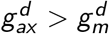 for mouse and human pyramidal cells, respectively; the divergence between the two species is due to the larger membrane resistance of human cells with respect to mouse’s cells, cfr. Table 2. **(B)** Effective membrane timescale 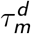 as a function of the dendritic length when the diameter is fixed at 2 μm (thin) or 4 μm (thick). Colors correspond to panel **A** and indicate the distinct functional regions of dendritic geometry. Thick dendrites influence somatic spiking more than thin ones, regardless of length. Mouse membrane timescale (dotted) converges with length while human timescales (solid) continue to increase. Throughout this work we will use the labels *proximal* and *distal* to refer to dendrites 150 μm and 400 μm long.

To test the sensitivity of the Tripod neuron to biophysical constraints, we compared two sets of membrane parameters corresponding to mouse (Koch, 1999; Dasika *et al*., 2007) and human (Eyal *et al*., 2016) layer 2/3 pyramidal cells. The axial conductance was the same across datasets, but the membrane conductance and capacitance differed (Table 3). To illustrate this difference, Fig. 3*A* shows the boundaries of effective dendrites in the Tripod neuron as a function of cable geometry. These boundaries correspond to the regions below which the membrane leakage is larger than the axial conductance, i.e., 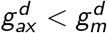. Consequently, dendritic currents fail to reach the soma in this case and the dendrite is rendered ineffective. Within our model constraints, the dendrites of human pyramidal cells can be substantially longer than those in mice and still be functionally effective, an observation that is consistent with recent empirical evidence (Fişek & Häusser, 2020). The functional role of dendrites is also dependent on the diameter of the dendritic compartment. Thin dendrites (2 μm) have low axial conductance and their contribution to the somatic voltage is small, i.e., thin dendrites are in the no-spiking region for most of their lengths. Thick dendrites (4 μm), on the other hand, place the neuron in the spiking regime for all the lengths considered.

### Human physiology supports longer dendrites

The effective membrane timescale characterizes the dynamics of the dendritic compartments. When the dendrite is depolarized and the soma is at the resting potential, the timescale 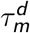 for the dendritic membrane to return to the resting potential depends on the physiological parameters. It is modulated by the dendritic length and diameter, as defined in Eq. 8. In the condition of effective dendritic transmission 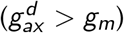, the current flowing out from the dendrites enters the somatic compartment, and the dendritic timescales together with the somatic membrane timescale fully determine the integration timescale in the Tripod model. Fig. 3*B* shows the integration timescale 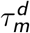 for all the considered dendritic lengths, two diameters (thin 2 μm, thick 4 μm), and the physiological parameters for human and mouse (solid and dashed lines, respectively). The membrane potential in longer dendrites decays slower because the axial conductance decreases and the capacitance increases with dendritic length. For a fixed diameter, doubling the dendritic length doubles the membrane timescale. Thin dendrites have a longer timescale because of the reduced membrane leakage and axial conductance.

Overall, the differences in membrane physiology and dendritic geometry constrain the membrane’s effective conductance and time constant and, consequently, the temporal integration properties of the neuron, leading to functionally relevant effects. Longer dendritic cables lead to sustained dendritic potentials, which affect the kinetics of somatic depolarization. This effect is particularly noticeable for human physiology, suggesting that human pyramidal cells can sustain longer functioning dendrites and that length modulates neuronal responsiveness significantly. Since the functional contribution of thin dendrites is limited, we focus on thick dendrites with a diameter equal to 4 μm, consistent with previous studies (Dasika *et al*., 2007; Yang *et al*., 2016; Bono & Clopath, 2017). In the remaining work, we will study dendritic lengths in the spiking region of the phase space, and this corresponds to dendrites in the range of 100 μm to 500 μm (blue and light blue regions in Fig. 3). For simplicity, we selected two lengths, 150 μm and 400 μm in the two spiking regions that satisfy Eq. 13 and Eq. 14. Following Antic *et al*. (2010) and Kamondi *et al*. (1998), we call a dendrite of 150 μm in length *proximal*, because it is capable of eliciting somatic spikes. A longer dendrite of 400 μm is referred to as *distal* and it can elicit somatic spikes only if co-activated with another dendrite. The proximal and distal dendrites described in the following sections are considered as roughly corresponding to the basal or apical oblique regions of pyramidal cells, respectively.

### Synaptic integration with segregated dendrites

The previous section investigated how dendritic geometry and membrane physiology determine temporal integration in the Tripod neuron. We now turn to the characteristics of synaptic transmission and how the existence of segregated dendritic compartments affects neuronal responses in the model. The synaptic models used are biophysically motivated and account for relevant physiological observations.

Due to their voltage-gated component, the dynamics of NMDA receptors (NMDRs) mediates the generation of sustained plateau potentials (Major *et al*., 2008) and supports coincidence detection (Tabone & Ramaswami, 2012; Rackham *et al*., 2010). It affects the integration of excitatory input in the dendrites and the soma, and plays a key role in shaping dendritic processing, synaptic plasticity, and the global input-output behavior of neurons (Doron *et al*., 2017; Smith *et al*., 2013). Furthermore, NMDAR expression is denser in distal regions along the dendrites (Larkum, 2013; Schiller *et al*., 2000) and this suggests that there is an important relationship between the geometry and the activation of the voltage-gated receptors.

We first investigated the influence of dendritic NMDARs on somatic depolarization and the magnitude of excitatory post-synaptic potentials (EPSPs). As explained in the Methods, we included NMDAR parametrizations corresponding to mouse (Avermann *et al*., 2012; Duarte & Morrison, 2019) and humans (Eyal *et al*., 2018). Compared to mouse, human NMDARs have shorter decay times, a larger NAR, and a steeper voltage dependence *γ* in the gating mechanism. In contrast, timescales and synaptic strength of AMPARs are approximately the same for the two species.

The experimental protocol used to test the effect of varying the NMDAR characteristics is shown in Fig. 4*A*. One of the segregated dendrites is stimulated with simultaneous spikes from excitatory presynaptic neurons, and the resulting EPSP is measured at the soma. In the synaptic model we used, coincident spikes corresponded to a single synaptic event whose efficacy was given by the peak conductance *g*_*syn*_ multiplied by the number of input spikes. The peak EPSP is identified as the difference in membrane potential between the moment of spike arrival and the maximal potential reached after the spike. The peak EPSP increases with the number of co-active presynaptic neurons and converges towards a maximum value determined by the axial conductance of the targeted dendritic compartment.

**Figure 4:**
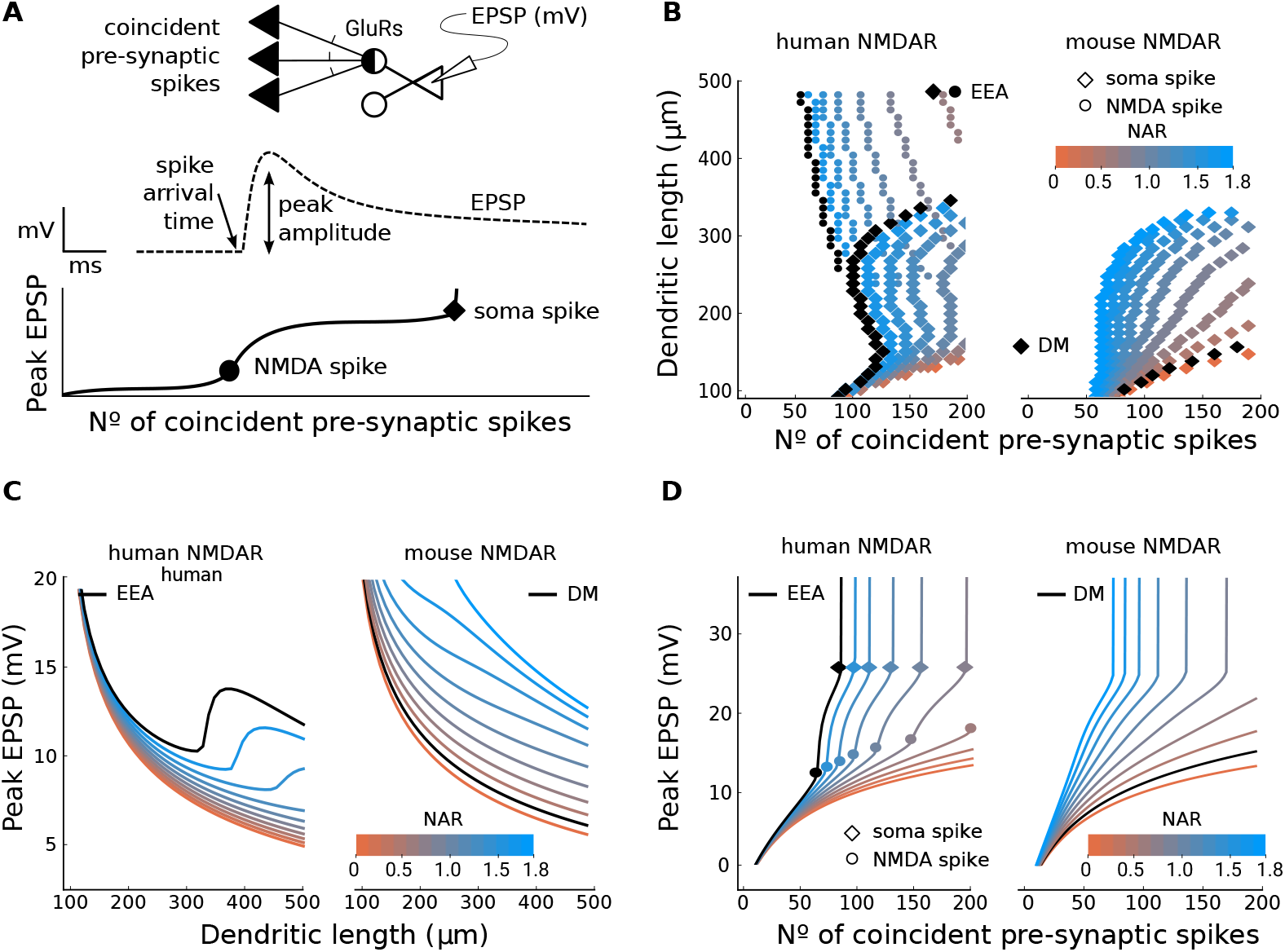
Human-like synapses induce NMDA-related supra-linearity in EPSP peak amplitude. **(A)** Schematic of the experimental setup. Multiple presynaptic spikes arrive concurrently at a segregated dendritic compartment with glutamatergic receptors (GluRs), and the resulting excitatory post-synaptic potential (EPSP) is measured at the soma (top). The peak amplitude of the EPSP is calculated as the difference between the membrane potential prior to stimulation and the peak membrane potential after stimulation (middle). Increasing the number of coincident presynaptic spikes results in larger peak amplitudes and causes NMDA spikes or somatic spikes (bottom). For an unbiased comparison of NMDARs between mouse and human parameters, the following simulations are based on human-like membrane parameters; when tested for mouse-like membrane, the EPSP response is weaker and sub-linear. **(B)** Tripod spike responses for *human* (left) and *mouse* NMDAR timescales and voltage gating slope (right). Each data point represents the minimal number of coincident presynaptic spikes necessary to elicit a somatic spike (diamond) or an NMDA spike (circle) for a given dendritic length (y-axis) and a specific ratio of NMDA-toAMPA receptors (NAR, color gradient). Note that NMDA spikes are absent for mouse synaptic physiology. Black markers show the spike responses for the combination of dendritic timescale and NAR described in Eyal *et al*. (2016) (labeled EEA) or Duarte & Morrison (2019) (labeled DM). **(C)** Peak amplitude of the EPSP as a function of dendritic length when the number of coincident presynaptic spikes is fixed at 60. *Human*-like synaptic parameters result in an upswing of the peak EPSP relative to the increasing dendritic length, which is weaker or absent for *mouse* parameters. **(D)** Peak amplitude of the EPSP as a function of the number of coincident presynaptic spikes when the dendritic length is fixed at 300 μm. While somatic spikes occur for both human and mouse NMDARs, only *human*-like synaptic parameters cause the supra-linearity in peak EPSP that is indicative of NMDA spikes (circles).

Segregated dendrites with NMDARs generate a supra-linear response in the somatic EPSP which is triggered when the dendritic membrane potential reaches the threshold of the voltagegated NMDARs. To track the onset of this supra-linearity, we computed the second derivative of the EPSP peak amplitude as a function of the coincident presynaptic spikes and determined its maximum. The onset is shown in Fig. 4*B* as a function of dendritic length and the number of coincident spikes. We distinguish between somatic spikes (peak amplitude of EPSP ≥ 30*mV*, diamond markers) and the NMDAR-related supra-linearity (circles). Because the opening of NMDARs causes an all-or-none event similar to the action potential, we also refer to the NMDAR supra-linearity as an NMDA spike. When glutamatergic synapses were parameterized according to human pyramidal cells (Eyal *et al*., 2018, Table 4), the NMDA-related non-linearity occurred alongside somatic spikes. When parameterized with a lower NAR, faster rise, and slower decay, corresponding to mouse synaptic physiology (Duarte & Morrison, 2019), the EPSP supra-linearity was absent, regardless of the number of synaptic inputs (Fig. 4*B*).

**Table 4:** Parameters for mouse (Duarte & Morrison, 2019) and human (Eyal *et al*., 2018) excitatory synapses. Inhibitory synapse parameters derived from Miles *et al*. (1996).

| Symbol | Description | Human |  | Mouse |  | Unit |
| --- | --- | --- | --- | --- | --- | --- |
|  | *Excitatory* | AMPA | NMDA | AMPA | NMDA |  |
| $E_r$ | Reversal potential | 0 | 0 | 0 | 0 | mV |
| $\tau_r$ | Rise time constant | 0.26 | 8 | 0.26 | 1 | ms |
| $\tau_d$ | Decay time constant | 2 | 35 | 2 | 100 | ms |
| $\bar{g}_{syn}$ | Peak conductance | 0.73 | 1.31 | 0.73 | 0.159 | nS |
| $\gamma$ | Voltage-gating slope | — | 0.075 | — | 0.062 | $\text{mV}^{-1}$ |

| Symbol | Description | Soma | Dendrites |  | Unit |
| --- | --- | --- | --- | --- | --- |
|  | *Inhibitory* | GABA<sub>A</sub> | GABA<sub>A</sub> | GABA<sub>B</sub> |  |
| $E_r$ | Reversal potential | $V_r$ | $V_r$ | -90 | mV |
| $\tau_r$ | Rise time constant | 0.5 | 4.8 | 30 | ms |
| $\tau_d$ | Decay time constant | 15 | 29 | 400 | ms |
| $g_{syn}$ | Peak conductance | 0.38 | 0.27 | 0.006 | nS |

The onset of NMDA spikes also depended on dendritic length. Fig. 4*C* shows a vertical section of Fig. 4*B* where the number of coincident spikes is fixed at 60 and dendritic length is varied between 100 μm and 500 μm. For mouse-like NMDARs, with fast rise and slow decay timescales, the peak EPSP decreased monotonically with the length of the dendrites. For human-like NMDARs, on the other hand, dendritic stimulation resulted in an increase of the peak EPSP amplitude for dendrites longer than 300 μm when the NAR was high. This indicates that the slow rise and fast decay timescales of human NMDARs and their higher voltage sensitivity were crucial in generating NMDA spikes. Fig. 4*D* is a horizontal section of Fig. 4*B* with dendritic length fixed at 300 μm. Somatic spikes occurred for both human and mouse NMDARs, but only human-like synaptic parameters caused the supra-linearity in peak EPSP that corresponds to an NMDA spike.

To summarize, the results suggest that a large NAR was not sufficient to elicit NMDA spikes in mouse-like NMDARs, regardless of dendritic length and the number of coincident presynaptic spikes. Increasing the NAR (Fig. 4*D*) raised the slope of the somatic response, but missed the supralinear component, which indicates that the supra-linear integration depends on the NMDAR steepness (*γ*) and timescales, which also differ between humans and mice (Eyal *et al*., 2018; Duarte & Morrison, 2019). For human-like NMDARs, the occurrence of NMDA spikes was mainly dependent on the NAR and dendritic length. The length of the dendritic compartment is a crucial variable for the rise of NMDA spikes; for the opening of voltage-gated ligands of NMDARs, the membrane potential has to be sufficiently depolarized (beyond ∼−40 mV). Such depolarization can happen only if the compartment is sufficiently electrically segregated from the soma and the other compartments, otherwise the membrane potential will leak towards the soma through axial currents. The dependence on dendritic length of NMDARs’ non-linearity confirms the importance of implementing voltage-dependent receptors in neuronal models with segregated dendrites

### Computation with minimal dendritic structure

The above results indicate that segregated compartments are necessary for the generation of NMDA spikes. However, models with a single dendritic compartment, usually referred to as ball-and-stick models, might not be sufficient to express important dendritic computations. For instance, several dendritic phenomena depend on the interaction among synapses and therefore on their spatial arrangement on the dendrites (Payeur *et al*., 2019; London & Häusser, 2005), and a cascade of synapses activated from distal to proximal sites elicits a stronger response than the reverse protocol (Branco & Häusser, 2010). Hence, the question is how many compartments are needed to express these computations? We argue that a minimal model requires two dendritic compartments because it can express a minimal form of dendritic branching and captures dendritic computations where the location of synaptic input matters. In a Y-branched dendritic tree, synaptic inputs can target the *same* or *different* dendritic branches, and the synaptic location becomes an important spatial variable of neuronal integration. This argument is in agreement with several *in-vitro* and *in-vivo* studies which have shown that two compartments are already sufficient to reproduce most of the observed processing complexity (Ujfalussy *et al*., 2018; Wybo *et al*., 2021). In the next sections we consider the Tripod neuron in three dendritic configurations, two symmetric (distal-distal and proximal-proximal) and one asymmetric (distal-proximal). We show that in the Tripod neuron the somatic response depends on the spatial location of the inputs and that two Y-branched dendrites are sufficient to express coincidence detection (Mel, 1992), inhibition-driven pathway selection (Yang *et al*., 2016) logical operations (Cazé *et al*., 2013). In addition, we introduce the concept of *dendritic memory* which is the neuron’s capacity to track previous activity in the voltage plateaus of distal dendrites. We show that dendritic memory can be utilized to integrate sequences of spatially distributed information and detect variations in the input stream.

### Coincidence detection

The conductance-based mechanism that transforms presynaptic events into currents and membrane depolarization determines the EPSP response to glutamatergic inputs that occur close in time. When two excitatory synapses fire together on the same dendritic branch, the combined effect can differ from two synapses firing on separate branches. For AMPA synapses, whose receptors are not voltage-dependent, synaptic inputs across spatially segregated dendrites is known to increase the somatic EPSP response, while clustered excitation on the same dendritic branch results in weaker EPSPs (Dasika *et al*., 2007; Li *et al*., 2019). The difference between clustered and spread inputs is caused by the interaction of conductance-based synapses with the compartment voltage (Koch, 1999). An increase in synaptic conductance produces weaker depolarizing currents if the compartment is already depolarized than if the compartment is close to the resting potential. A formal derivation of this interaction is provided in Appendix B. However, as demonstrated by Mel (1992), the expression of dendritic NMDARs can yield the opposite effect. For these receptors, clustered excitation can result in larger somatic EPSPs than spread excitation, which can be interpreted as a dendritic mechanism for coincidence detection.

To test whether the Tripod neuron can reproduce these clustering effects, we compared the EPSPs generated at the soma in two conditions, clustered and spread synaptic input, and tested how the spatial distribution of the input affected somatic EPSP responses. The Tripod model is investigated here with two symmetric dendritic compartments, labeled A and B. We used dendritic lengths of 150 μm and 400 μm which are representative of compartments with weak and strong segregation from the soma. These two configurations are referred to as *proximal-proximal* and *distal-distal* configurations. We measured the difference ΔEPSP between the somatic EPSPs resulting from excitatory input that was spread over the two compartments (EPSP_AB_) or clustered on one compartment (EPSP_AA_) as shown in Fig. 5*A*. Negative values for ΔEPSP indicate that the global synaptic current was reduced for clustered input relative to spread input, whereas positive values indicate that the somatic peak depolarization was stronger for clustered input relative to spread input. ΔEPSP was measured for 200 simulations, with a random number of co-active synapses drawn uniformly from the interval [1, 50] for each branch A and B in order to simulate different input intensities. The results are shown in Fig. 5*A* where the x-axis shows the total number of co-active synapses on the two branches. There was no difference between proximal dendrites that expressed NMDARs or AMPARs only. In both cases, input spread across dendritic branches generated a larger somatic EPSP than clustered input, and this was also the case for distal dendrites with AMPARs only. However, for distal dendrites that also expressed NMDARs, clustered input caused a larger EPSP when the total synaptic input was strong, as indicated by the positive ΔEPSP (orange data points) in Fig. 5*A* (bottom left). Thus, the Tripod neuron reproduces the AMPA spread effect and the NMDA clustering effect described in the literature (Mel, 1992; Dasika *et al*., 2007).

**Figure 5:**
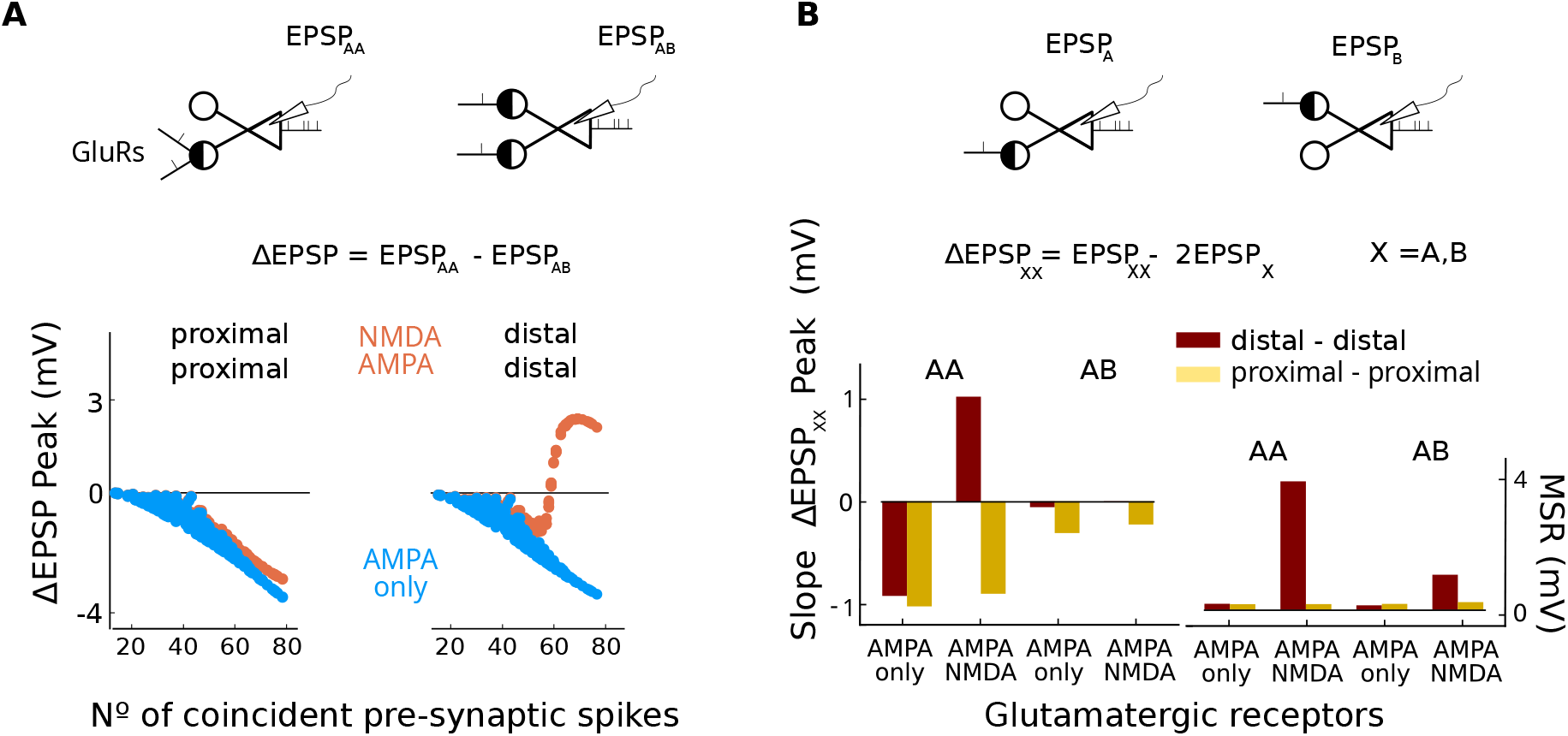
NMDA receptors enhance somatic response in clustered condition. **(A)** Excitatory input was applied on one dendritic branch only (clustered, AA), or on both dendritic branches (spread, AB), and the elicited EPSPs were measured at the soma. The difference between EPSPs in the two conditions is denoted ΔEPSP (top panel). The two dendritic branches had the same length and were either distal or proximal. Synapses on the two branches expressed NMDA and AMPA receptors (orange), or AMPARs only (blue). The bottom panel shows the peak ΔEPSP as a function of the number of coincident input spikes in the four conditions. For proximal-proximal dendrites, spread input resulted in stronger EPSPs for both AMPARs only and combined AMPARs/NMDARs. For distal-distal dendrites, the expression of NMDARs produced stronger responses in the clustered condition which showed a supra-linear response when the total synaptic input was sufficiently strong to activate the NMDARs (>60 co-active synapses). **(B)** The magnitude of synaptic interaction was obtained by comparing di-synaptic conditions (*XX* = AA, AB) to input spikes on single compartments (*X* = A, B). The top panel shows the stimulation protocol used to compute EPSP_A_ and EPSP_B_. ΔEPSP_AA_ and ΔEPSP_AB_ summarize the interaction for the clustered and spread conditions. The ΔEPSPs in conditions AA and AB are fit with linear regression over the global synaptic inputs and the lower panels show the slope and the mean squared residuals (MSR) of the linear fit. Di-synaptic interaction reduced somatic depolarization (negative slope of ΔEPSP_AA,AB_) for all inputs conditions, receptor types, and Tripod configurations except for clustered inputs AA on distal-distal compartments with NMDARs (third column). This configuration generated high MSRs, indicating that the interaction could not be expressed with linear di-synaptic interactions. For all conditions the fit was computed by drawing 200 co-active synapses in the range (1,35).

To disentangle the effects of physiology and geometry, we attempted to estimate the nonlinearity of the EPSP-response based on the second-order model proposed by Li *et al*. (2019). The original model introduced a di-synaptic matrix *α*_*ij*_ that determines the difference in synaptic current with respect to two synapses firing independently. The values of *α*_*ij*_ depend on the efficacy and the location of the synapses that are active simultaneously. They are small for synapses on different branches, and negative for synapses on the same branch. To demonstrate that this second-order model is not sufficient to explain the synaptic interaction in the presence of voltage-dependent receptors in segregated dendritic compartments, we stimulated the Tripod with clustered and distributed inputs and subtracted the EPSP of independent synaptic event on each branches

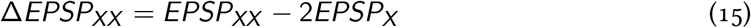

where the *X* subscript refers to the branches A or B. Note that EPSP_A_ is the same as EPSP_B_ because dendrites were symmetric. ΔEPSP_AX_ was computed for different numbers of co-active synapses between 1 and 50, as before. The simulation was run for 8 conditions, i.e., with and without NMDARs, with two distinct geometries, and in both the distributed AB and clustered AA configurations. Following the original model, we asked whether a second-order function of the synaptic input was sufficient to explain ΔEPSP_AB_. Hence, we fit ΔEPSP_AB_ via the product of the synaptic conductances 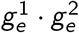 and obtained the results in Fig. 5*B*. The two panels show the slope of the interaction corresponding to *α*_*ij*_ in Eq. 23 and the residuals of the linear fit (right).

In the absence of NMDARs, we observed a strong attenuation of somatic EPSPs and the residuals of the linear fit were small. This effect was larger when synapses clustered on the same compartment compared to the distributed condition and this was due to the segregation of voltages in the different compartments. The EPSP attenuation effect was also stronger when dendrites were shorter (proximal-proximal configuration, yellow bars in Fig. 5*B*). It is worth noting that the residuals of the linear fit were small for most of the configurations, suggesting that the model of Li *et al*. (2019) was also applicable to the Tripod neuron when only AMPA receptors were present. However, in agreement with previous results (Mel, 1992), the expression of dendritic NMDARs yielded different functional behavior and resulted in the amplification of somatic EPSPs in the clustered condition AA. This effect was dependent on dendritic length. The di-synaptic interaction still resulted in EPSP attenuation (negative) in the proximal-proximal configuration due to the reduced NMDAR contribution for proximal dendrites. For longer dendritic branches (distal-distal configurations in Fig. 5*A*), when excitatory inputs were clustered on the same compartment, the interaction initially reduced somatic EPSP amplitudes. As the number of co-active synapses increased to around 60, however, the EPSP began to increase in a non-linear fashion. Thus, segregated dendritic compartments with voltage-dependent NMDA receptors introduce synaptic interactions that go beyond the second-order model of Li *et al*. (2019). These interactions cause larger EPSPs when synaptic inputs are clustered, in agreement with previous simulations (Ujfalussy & Makara, 2020; Mel, 1992), and the magnitude of this clustering effect is strongly mediated by dendritic length.

### On-path shunting inhibition

Depending on the location of synaptic contact, inhibitory GABAergic inputs, whose ionotropic receptors have an equilibrium potential close to the resting potential, can effectively offset excitatory drive onto neighboring synapses (Koch *et al*., 1983). Inhibitory configurations that veto neuronal responses are referred to as *shunting* inhibition and play an important functional role. Shunting inhibition depends on the spatial distribution, the composition of inhibitory synapses, and the relative timing between excitatory and inhibitory presynaptic events. The Tripod neuron with two dendritic and one somatic compartment provides the simplest structure to study this type of inhibition.

We investigated different inhibitory configurations by stimulating one of the dendritic compartments with a single excitatory spike followed by an inhibitory spike within a fixed time interval that was delivered to one of three input locations; the same dendritic compartment (*on-path*), the other dendritic compartment (*off-path*), or the soma (Fig. 6*A*). To measure the effectiveness of inhibition, we compared the somatic EPSP in the presence or absence of GABAergic inputs. Attenuation caused by inhibition was measured as the ratio between the EPSP peaks in the two protocols (excitation versus excitation plus inhibition):

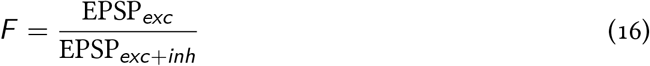

The larger this *F* -factor, the more effective the inhibitory signal was.

**Figure 6:**
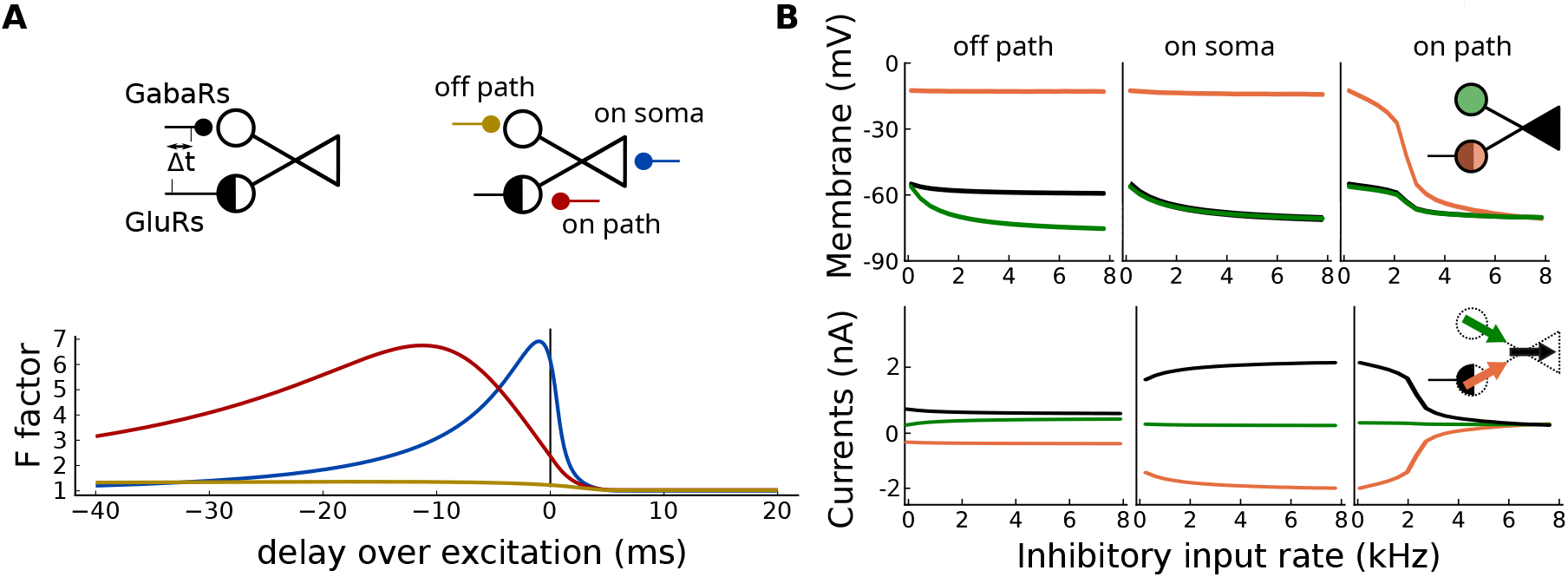
Dendritic inhibition and shunting. **(A)** Location and timing of inhibitory spikes determine the somatic response. The upper panel describes the experimental protocol. An inhibitory spike is delivered to the soma at an interval Δ*t* from the excitatory one. Then the F-factor is computed. The two panels show the EPSP attenuation for three inhibitory conditions, on-path (red), off-path (yellow), on soma (blue), scheme in the lower panel. In the upper panel, the dendritic GABA_A_ receptors are parameterized with long timescales, as in Miles *et al*. (1996). **(B)** Average membrane potentials (upper panel) and axial currents (lower panel) for varying inhibitory input rates. The orange dendrite receives 1.5 kHz Poisson distributed excitatory input, while the neuron also receivs variable inhibitory inputs at different locations (from the left: off-path, on-soma, on-path). Both dendrites are 300 μm long. Inhibition off-path has a negligible effect on the somatic membrane (black line) compared to on-path and on-soma inhibition.

Results in Fig. 6*A* show that the impact of inhibition is determined by the relative timing of the excitatory and inhibitory inputs and it is highly location-specific. Suppose dendritic GABAergic transmission in the same compartment of excitation, *on path*. In that case, its depressing effect on the EPSP is extended in time, and it peaks when inhibitory spikes arrive around 10 ms before excitation (red line). If, on the other hand, inhibition is located on the soma, hence mediated by fast GABA_A_ receptors, then inhibition is maximally effective when inhibitory and excitatory inputs arrive simultaneously. In this condition, inhibitory spikes that arrived more than 10 ms before excitation are ineffective. When inhibition is *off-path* its effect on the somatic EPSP is negligible. Notice that the GABA_B_ receptors are active only in the dendrites, and their effect is small in the setup of Fig. 6*A* because a single inhibitory spike is insufficient to engage these receptors.

The Tripod neuron received an excitatory Poisson input at a fixed rate of 1kHz on a single dendritic compartment and a variable rate inhibitory input on different compartments (off-path, on soma, on-path). In the absence of inhibitory input, the soma was in a depolarized state 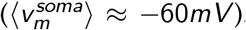. Fig. 6*B* shows the mean value of the membrane potential of each compartment and the current flowing between the compartments, both averaged over a 10 s interval. When the *off-path* compartment was targeted by inhibitory inputs (leftmost panel in Fig. 6*B*), the soma reached equilibrium between a weak hyperpolarizing current coming from the inhibited dendrite and the depolarizing current from the excited compartment. In this condition, the soma remained depolarized, regardless of the magnitude of the inhibitory inputs. When inhibition targeted the somatic compartment (middle panels in Fig. 6*B*), the soma received a depolarizing current from the excited dendrite and a competing hyperpolarizing current from the GABAergic synapses on the soma membrane. Because the synaptic current depended on the somatic potential, it had a balancing effect on the compartment potential. When the inhibitory input was sufficiently strong, the soma approached the resting membrane potential. In this condition, inhibition had a divisive effect on the somatic potential. In both cases, the stimulated dendrite remained depolarized but benefitted from the NMDA boost, resulting in a large axial current. On the other hand, when the inhibition was *on-path*, that is, localized to the same compartment as excitation, inhibition pulled the dendritic potential below the NMDA threshold, and thus hyperpolarized the stimulated dendritic compartment. In this configuration, the soma remained depolarized as long as the dendritic balance of excitation and inhibition was maintained. When the inhibition overcomes excitation (around 2 kHz for this setup), the neuron was shut down, and all the compartments went to resting potential, with no axial currents flowing. Hence, somatic depolarization is more dependent on the spatial distribution of the inhibitory spikes than on the actual inhibitory input received. Furthermore, this experiment suggests that considering the somatic membrane potential alone may not be sufficient to characterize the state of the cell; in Fig. 6*B* the membrane potential of *on soma* and *on path* conditions is similar, although the cell is in two different states and will respond differently to further stimuli. For example, for a fixed inhibitory input, increased excitation on the stimulated dendrite will only depolarize the soma if inhibition is *on path*, while it will be less effective in the *on soma* condition

### Logical operators

Logical operators define a natural class of computations. Single compartment neurons, which integrate inputs with a monotonic transfer function, can perform linearly separable computations but fail on non-separable ones. In contrast, theoretical and experimental work has shown that active dendrites can solve non-separable problems (Cazé *et al*., 2013; Gidon *et al*., 2020). If we consider the dendrites as independent input pathways and treat the Tripod as a binary logical gate, then the previous experiments on coincidence detection have already demonstrated that the Tripod can perform non-separable computations, matching the theoretical results in (Cazé *et al*., 2013).

Another possibility is to consider the neuron’s dynamics explicitly. In this configuration, the input is drawn from a set of binary stimuli, e.g., A = 0, B = 1, and mapped to the input spike rates on the respective compartment, e.g., 0 = E/I balanced inactive state, 1 = E/I balanced active state (further details in Appendix C). The cell’s response also has to be represented over time and calculated, for example, on the output firing rate. Under this encoding, both a single-compartment neuron and the Tripod model can reproduce the truth table of multiplication (AND, true for inputs (1, 1)) and summation (OR, true for inputs (0, 1), (1, 0), (1, 1)). However, there are no mechanisms that enable single neurons to implement operators such as exclusive OR (XOR, true for (1, 0) and (0, 1) but false for (1, 1) and (0, 0)) or material implication (MI, true for (0, 1) but false for (1, 0)). Unfortunately, the same holds for dendrites with NMDA spikes; if one active dendrite is sufficient to trigger somatic spikes, two active dendrites can only increase the somatic firing rate, making it impossible to solve the XOR problem. These limitations are due to the coding scheme for the output. To avoid this, we investigated if the neuron could make the computation separable for an external linear readout. Therefore, we analyzed the sub-threshold dynamics of the somatic membrane potential (van den Broek *et al*., 2017) to evaluate the neural computations.

For this purpose, we stimulated the dendrites with a random sequence of four possible input configurations: (A = 0, B = 0), (A = 1, B = 0), (A = 0, B = 1) and (A = 1, B = 1). A set of seven external logistic regression readouts were used to map the neurons’ somatic dynamics to the truth table of seven different operators (*Id*_*A*_, *Id*_*B*_, A ∨ B, A ∧ B, A ⊕ B, A ⇒ B, B ⇒ A) As mentioned above, the symbols A and B refer to the stimulated dendritic compartment and each input is presented for a period of 200ms. The membrane dynamics was readout during the last 50 ms of the stimulus presentation; the readout had access to 5 points for the membrane potential and 5 points for the adaptive current, each spaced by 10 ms. After training, we injected a random sequence of inputs (A, B, AB, or none) and tested if the trained readout could use the information in the membrane of the soma to reproduce the correct truth table. We examined four different geometries, two symmetric one with proximal-proximal (150 μm) or distal-distal (400 μm) dendrites, one with asymmetric structure (400 μm-150 μm) and a single-compartment model. When a dendritic pathway was inactive (e.g., A = 0), the respective dendrite received a 3 kHz train of excitatory Poisson spikes, and a balanced inhibitory input. For the baseline condition (soma-only), the spikes were injected into the somatic compartment via two independent synapses, as above, the excitatory input rate was doubled for the active input condition.

After testing all the models, we measured the Coheh’s kappa-score of the readout on each operator, see Fig. 7*A*; we chose this metric to account for asymmetries in the classes’ statistics, e.g., A ⇒ B has three True and one False. Symmetric configurations performed better on symmetric operators (blue bars in AND, OR and XOR operators). Conversely, asymmetric operators (red bar in *Id*_*B*_, A ⇒ B) are best recognized with asymmetric dendrites. In the distal-proximal configuration, the activity in each dendrite is different, and input to the short dendrites is easily distinguished. The soma-only configuration scores lower than each Tripod configuration. To elucidate the computations performed, we analysed the predicted truth-value for each operator and condition (Fig. 7*B*). As expected, the symmetric configuration (proximal-proximal, distal-distal, and soma-only) makes the same prediction concerning inputs (1, 0) and (0, 1); for asymmetric operators, this is also the case, because the readout cannot distinguish which input-pathway is activated. This is not the case for the proximal-distal condition, and the input (0, 1) is treated differently from (1, 0). In almost all conditions the Tripod neuron performed better than the single-compartment model, indicating that the inclusion of the dendritic structure was beneficial. These results show that the membrane dynamics of asymmetric Tripod models depends on the input pathway, and the neuron can act as an asymmetric logical operator.

**Figure 7:**
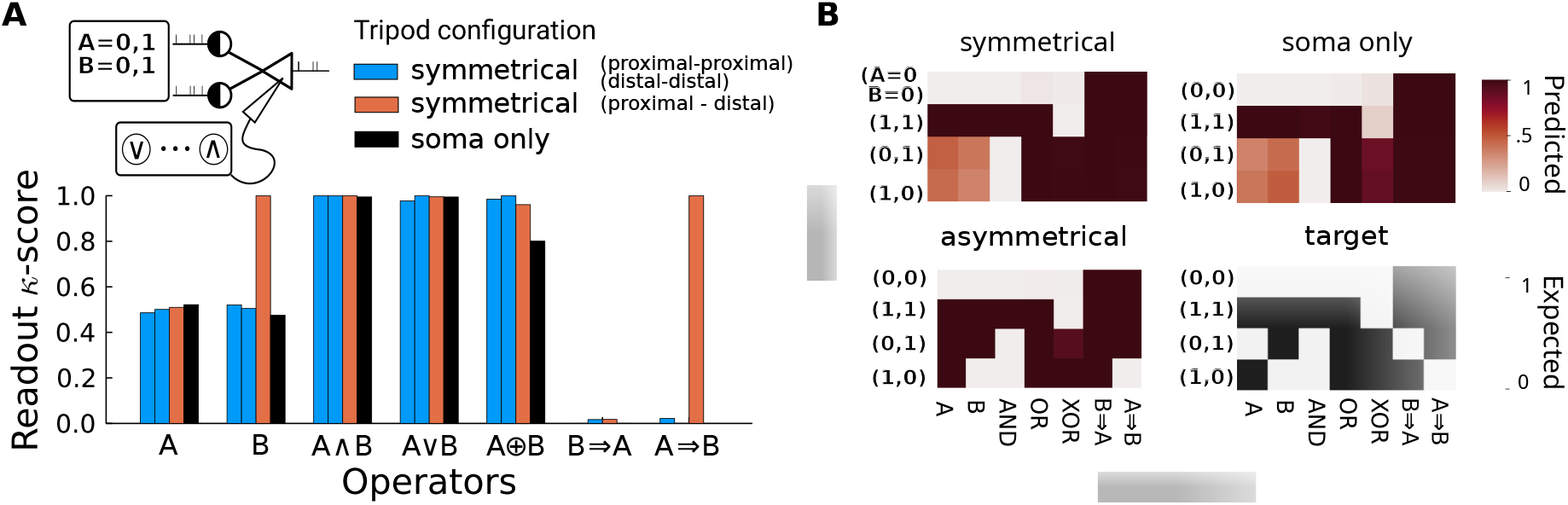
Asymmetric dendrites enhance separability of logical operations. **(A)** Cohen’s kappa-score accuracy of linear readout classifiers on logical operators for symmetric, asymmetric and soma-only models. The dendritic configurations are proximal-proximal and distal-distal (blue), proximal-distal (orange) and soma-only (black). **(B)** Shade of red indicates the average predicted truth-value for each input condition (y-axis), operator (x-axis), and dendritic configuration (top and left panels). Black and white table (bottom-right) indicates the expected truth-values. E.g., the AND operator for symmetric dendrites shows dark red (true) for condition A = 1, B = 1, and white for all the remaining conditions, corresponding to the target truth-values.

### Dendritic memory

When excitatory synaptic input is sufficiently strong to drive the postsynaptic voltage above the NMDA gating threshold, the ionic current flowing through the NMDAR keeps the dendritic compartment depolarized and generates a temporally extended plateau potential (Fig. 8*A*).

**Figure 8:**
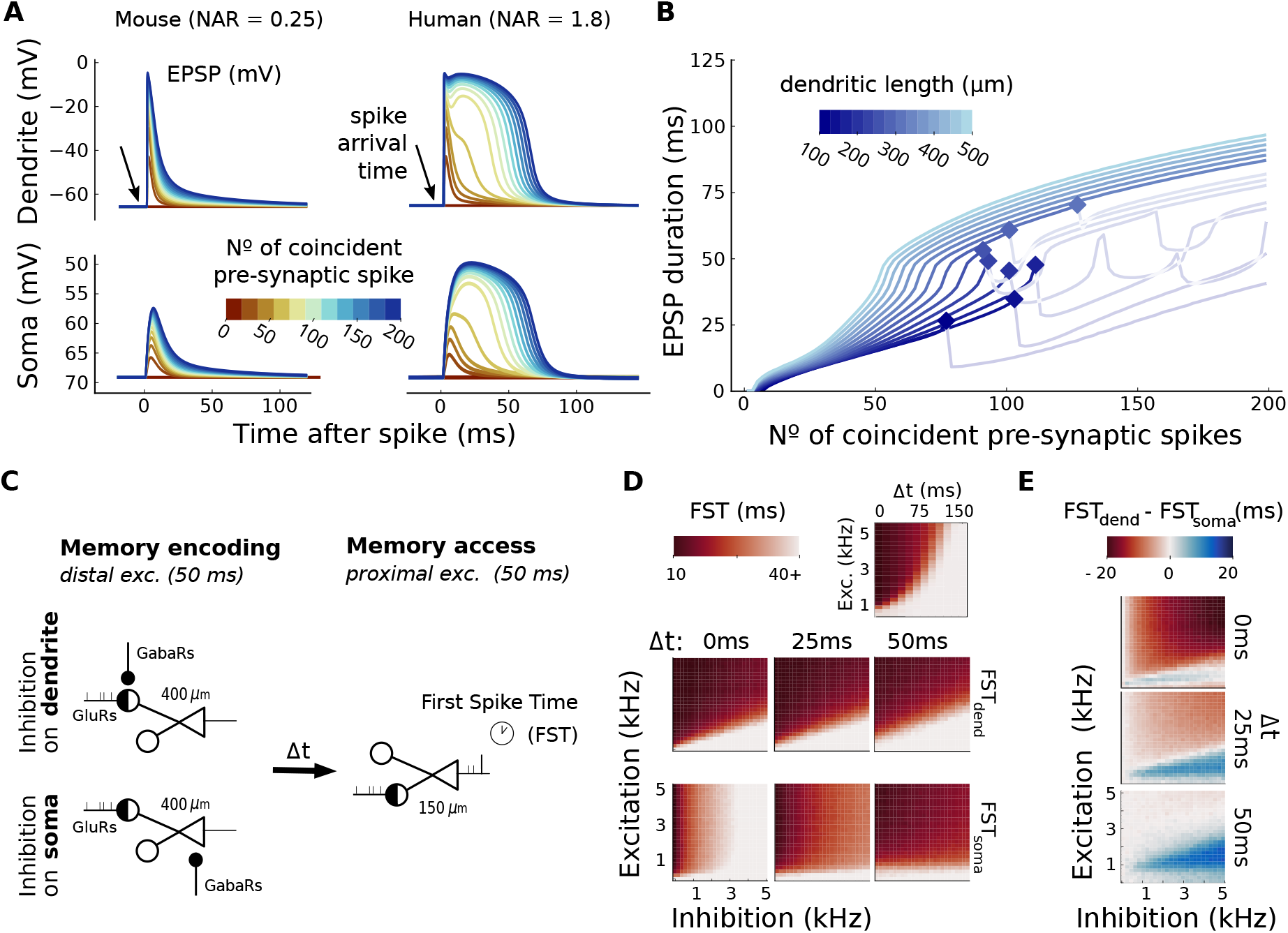
Plateau potentials due to NMDA spikes support dendritic memory. **(A)** Magnitude and kinetics of spike-induced EPSPs in the dendrites (upper panels) and soma (lower). Dendritic synapses are endowed with mouse (left panels) or human (right) NMDARs; AMPARs are identical for both. Input spikes arrive on a distal dendrite (400 μm), color codes for the number of coincident input spikes. For mouse-like synapses, an increase in the number of inputs did not lead to longer dendritic depolarization. Dendrites with human NMDARs show extended depolarization when the input triggers NMDA spikes. **(B)** Duration of sustained somatic depolarization (EPSP curve is above -60 mV) for simulations with *human*-like NMDARs. Color codes for dendritic length. Long dendrites result in a somatic depolarization that lasts for +100 ms, referred to as *plateau potential*. For long dendrites, the duration of the plateau potential increases monotonically with the number of simultaneous synaptic inputs. When the targeted dendrite is short enough to cause somatic spikes (diamond markers), the relation between the total presynaptic input and the duration of the depolarized state is interrupted because the somatic after-spike depolarization forces the dendrite below the activation threshold of the NMDARs. Somatic spikes do not affect the plateau potential in long dendrites because of the low axial conductance. **(C)** Input configuration for memory encoding. Memories are encoded via excitation of the distal dendrite. After an interval Δ*t* without input, the proximal compartment is activated and the average first-spike-time (FST) is measured. Figure 8: **(D)** Mean FST for varying excitation strength and Δ*t* (upper-right). Shorter FSTs (e.g., dark red, 10 ms) indicate successful memory retrieval. Lower panels show FSTs for dendritic (upper) or somatic (lower) inhibition, measured while varying the strength of excitation and inhibition during the encoding phase. Retrieval is attempted after three intervals Δ*t* ∈ {0, 25, 50} ms. **(E)** Comparison of memory traces in two inhibitory configurations. Colors code for the difference between FSTs in the somatic versus dendritic inhibition condition. For short Δ*t*, inhibition on dendrites elicits faster somatic spikes (shorter FST). For longer Δ*t*, inhibition on dendrites is more detrimental to retrieval than inhibition on soma.

The time-course of the plateau potential depends on the number of coincident presynaptic spikes, even though the dendritic potential reaches the NMDAR reversal potential (Fig. 8*A* top right panel). To quantify the duration of the voltage plateau, we set an arbitrary threshold at −60 mV and monitored how long the somatic membrane potential remained above this value (Fig. 8*B*). In the presence of NMDARs with *human* timescales, NAR, and *γ*, long distal dendrites reached a voltage plateau whose duration increased with the number of coincident inputs and could last up to 100 ms. When dendritic length was short enough to trigger somatic spikes (Eq. 13) the duration of the plateau potential was limited by the somatic after-spike reset potential. Because of the large conductance between proximal and somatic compartments the brief duration (1 ms) of the hyperpolarized reset potential is sufficient to prevent the continuation of the plateau-potential by pulling the dendritic potential below the NMDAR threshold. Conversely, this is not the case for distal dendrites that can sustain the plateu-potential during somatic firing. Further details on dendritic membrane dynamics during and after somatic spikes are discussed in Appendix C. When the NAR was set to mouse synapses (0.25), the dendritic and somatic potentials showed a weaker, sub-linear dependence on the number of presynaptic inputs. The depolarization caused by 50 synapses was similar in extent to the depolarization caused by four times as many coactive synapses (Fig. 8*A*, left panel). The reason why the EPSP response saturates is because the incoming synaptic current depends on the difference between the membrane potential and the synaptic reversal potential. For the remainder of this article, the dendritic parameters were set to correspond to human physiology.

We investigated whether the plateau potential generated by NMDA spikes in distal dendrites could be used as a short-term processing memory. We tested this by encoding a memory trace into distal dendrites through synaptic activity. The spike rate of the encoding signal was the critical variable and corresponded to the number of co-active synapses in the previous experiment. Then, we attempted to retrieve this memory by injecting a 1 kHz spike train on the proximal dendrite after an interval of time Δ*t* (illustrations in Fig. 8*C*). The retrieval cue was weak and without previous distal inputs the soma fired the first spike on average 50 ms after the onset of the proximal input. Note that the proximal input lasted longer than the 50 ms considered for retrieval. Thus, we considered retrieval of an encoded memory to be successful if the first somatic spike occurred earlier than 40 ms after the retrieval cue was injected. This measure of retrieval was called the first-spike-time (FST) and averaged over 300 independent trials in the experiment. The somatic compartment was also exposed to noisy excitatory inputs that caused random spikes during the stimulation protocol. This was not necessary for encoding and retrieval but was intended to test the robustness of plateau potentials in the presence of somatic spikes. The top-right panel in Fig. 8*D* shows that memories encoded into long dendrites could be retrieved within about a hundred milliseconds, which was approximately the duration of the plateau potential. The lifetime of memory traces increased with the input rate that was used to encode these memories (y-axes). However, higher input rates during encoding did not correspond to shorter FSTs.

So far, only glutamatergic synapses were considered. We further investigated dendritic memory in the presence of inhibition by activating GABAergic synapses during and right after the encoding phase. Inhibition was present on both somatic and distal compartments. We tested memory retrieval by monitoring the FST at three different times, separated by 25 ms each, after the encoding phase. Fig. 8*D* shows the effect of inhibition on the distal dendrite and on the soma. When excitatory inputs on the distal dendrite were matched by dendritic inhibition, retrieval depended on the ratio between excitation and inhibition, as demonstrated by the linear separation between successful and failed retrieval. The retrieval protocol cannot distinguish between the exact amount of inhibition received during the encoding phase when memory was successfully encoded; the upper panels in Fig. 8*D* show nearly identical success rates in memory access for the three delay intervals. This changed when inhibitory synapses fired on the soma; at first, memories were not retrievable but they became accessible when inhibitory activity ceased. Within 50 ms, there was virtually no trace of the somatic inhibition. In this condition the magnitude of the inhibitory input modulated the retrieval success rate in a graded manner.

The difference between the two inhibitory input pathways is shown in Fig. 8*E*. Immediately after distal activity (Δ*t* =0 ms), inhibition on the soma prevented spiking and memory retrieval (FST with dendritic inhibition was smaller than FST with somatic inhibition, dark red). After 50 ms, the relation between somatic and dendritic inhibition reversed and memories that were encoded during somatic inhibition were now accessible. Dendritic inhibition limited the life-span of the encoded memories and the ratio between excitation and inhibition during the encoding phase determined retrieval success. This shows that the minimal dendritic tree of the Tripod model maintained short-lived memories. Retrieval of these memories depended on the location, the input strength, and the relative timing of their encoding.

### Transition detection and sequence recognition

Dendritic memory endows the Tripod model with two segregated memory slots, which can potentially be used to combine or discriminate incoming information over time. Here we tested whether this memory mechanism could be used to solve spatio-temporal tasks.

Excitatory and inhibitory Poissonian inputs were injected into the neuron at a constant rate. The active dendrite was set in the E/I balanced active state, the other dendrite in the inactive state (further details in Appendix C). The input targeted dendrite A or dendrite B and it was switched from one compartment to the other regularly, with frequencies in the range of 1 Hz to 100 Hz. A schematic of the input protocol is shown in Fig. 9*A*. We first measured dendritic and somatic potentials during a sequence of switches at 4 Hz. The membrane dynamics of the three compartments is shown in Fig. 9*B*, for models with symmetric dendrites (distal–distal) asymmetric ones (distal–proximal). The sequence of excitatory and inhibitory input spikes was the same for the two models. After a switch, the potential of distal dendrites decayed slowly while the potential at the newly stimulated dendrite started to rise. As a consequence, the depolarizing axial currents towards the soma reached their maximum right after the switch. To measure the effect of the increased axial currents we computed the average somatic potential for 300 trials with similar input statistics (Fig. 9*B*). The somatic response to a switch differed between the two dendritic configurations. For distal–distal dendrites the response was maximal right after the switch and it was the same for the two dendrites. For distal–proximal dendrites, the somatic response was stronger during stimulation of the proximal dendrite than the distal dendrite, resulting in somatic bursts.

**Figure 9:**
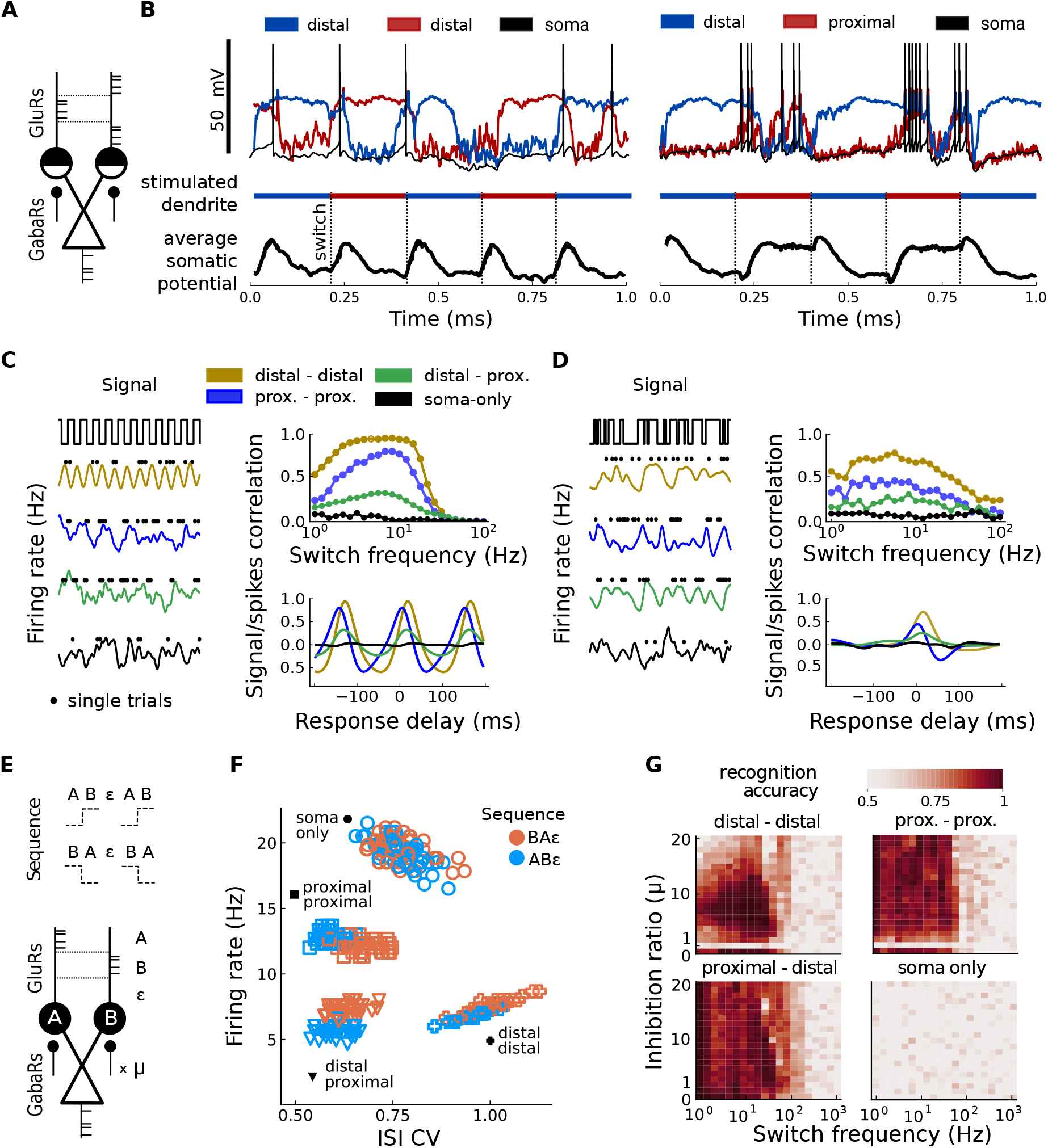
Sensitivity to serial order. **(A)** Excitatory and inhibitory inputs are delivered to the neuron by switching between the two dendrites periodically after a fixed interval. **(B)** Distal– distal and distal–proximal Tripod neurons receive the input described in (A). Each dendrite depolarizes during its stimulation interval. For distal dendrites, decay to rest is slow and the depolarized state overlaps in time with the rise in potential of the other compartment. This overlap of the two depolarized dendritic states maximally depolarizes the soma, as shown in the average membrane potential of the somatic compartment (lower panels). For asymmetric dendrites, somatic depolarization is strong when the proximal compartment is stimulated. Input to the two dendrites switches at 4 Hz and the average over 300 trials is shown. **(C)** Left panel shows average firing rate in response to a signal switching between dendrites at (6 Hz) for three Tripod configurations (colors) and a single-compartment model (black) that implements switching on two independent synaptic conductances. Black dots show spike-times on one of the 300 trials used to compute the firing rate (solid lines). Top-right panel shows the correlation between firing rate and switches in the input signal as a function of switch frequency. To compute the correlation, the switch times were convolved with an alpha-function. Bottom-right panel shows the correlation when the firing response is shifted in time (backward or forward) for inputs with 6 Hz switch frequency. The correlation is maximal after a delay for all the models because the soma lags behind the dendritic depolarization. Values shown in the top panel correspond to the maximal correlation obtained across all the response delays that were tested. Negative delays are due to the convolution function that maps spikes to rates. Peaks at ±150 ms are due to the oscillatory nature of the input. **(D)** As in (C) for a signal whose switch times are drawn from an exponential distribution of rate equal to the switch frequency. **(E)** Two sequences are played to the dendrites A and B of the neuron, AB*ϵ* or BA*ϵ*, where *ϵ* is a silent pause. Dendrites receive feedback inhibition proportional to somatic activity. One of the dendrites receives *β* times the feedback inhibition of the other dendrite. **(F)** With *β* β= 1, the spike statistics (firing rate and ISI CV) depend on both the sequence order (blue or orange) and the neuron’s geometric properties (marker shapes). Each data point corresponds to 1 s of simulation time and the switch frequency was 6 Hz. **(G)** Sequence classification accuracy based on the somatic spike statistics in (F) as a function of inhibitory feedback ratio *β* and switch frequency. Neuron configurations with dendrites outperform a soma-only model. Only the asymmetric configuration succeeded on the task when inhibitory feedback was identical on both dendrites (*β* = 1).

To further explore the Tripod’s response to spatio-temporal sequences, we tested four dendritic configurations, distal-distal, proximal-proximal, distal-proximal and soma-only. For a fair comparison with the soma-only model, the switching was achieved by implementing two independent synapses that were targeted by one of the two input streams. This corresponds to a model with zero-length dendrites. We repeated the previous experiment with two input signals, one with regular switching times as above, and one where switching times were drawn from an exponential distribution with rate equal to the switching frequency. We recorded somatic firing and averaged the output spikes over 300 trials with identical statistical realizations of the input spike train. Therefore, the reported firing rates indicate the average instantaneous somatic firing. The firing rate in response to the two input signals is shown in the left panels of Fig. 9*C* and *D*. Somatic firing on single trials was not synchronized with the switch times (black dots), but across multiple trials it was. To quantify the dependence of output firing on the input switch, we convolved the switch times with an alpha-function (with rise and decay timescales of 10 ms) and then computed the correlation between the average firing rate and the distribution of switch times (referred to a signal/spikes correlation in the top right panels of Fig. 9*C* and Fig. 9*D*). Overall, the Tripod responses were correlated to the spatial switches in the input stream, for both regular and irregular switching times. As expected, the distal–distal model showed the strongest correlation with the input switch, and the peak firing rate of a Tripod with asymmetric dendrites was less synchronized. For regular switch intervals the Tripod model lost track of signals oscillating faster than 30 Hz, although the response to non-regular signals stayed synchronized for higher frequencies. For both input conditions, the soma-only model showed zero correlation with the switching times. As suggested by the delays between the switch times and the maximal somatic response in Fig. 9*B*, we hypothesized that the correlation might be higher at different time points. Hence, we measured the correlation backward and forward in time with delays in the range of −200 ms to 200 ms. The correlation with the signal was maximal when the firing response was correlated backwards in time with as the somatic response lagged behind the input signal. The optimal delay depended on the model, and it was shorter for shorter dendrites. The bottom panels of (Fig. 9*C* and *D* bottom panels) show the correlation for different delays and an input signal with 6 Hz switch frequency. The delay with the highest correlation was the same across all the switch frequencies tested (data not shown), indicating that the optimal delay depended only on the time-span necessary to depolarize the dendritic compartment, which in turn depended on dendritic length.

These results show that the Tripod neuron with symmetric dendrites was sensitive to transitions in the location of synaptic input. We further investigated whether the switching *direction* could be detected as well. Two input sequences were created where input was injected into dendrite A, then dendrite B, or the other way around, followed by an inputless pause *ϵ* (see Fig. 9*E*), resulting in two sequences *ABϵ* and *BAϵ*. The switch intervals were regular and we use the switch frequency to indicate the rate for rotating over the elements of the sequence (A,B,*ϵ*). As before, the input spike trains targeting each dendrite were statistically the same. In a preliminary analysis we measured the somatic potential during presentation of the two sequences and verified that for symmetric models it was impossible to determine which of the two was presented. When the model had asymmetric dendrites, the order of the input on the dendrites (proximal-distal-*ϵ* vs. distal-proximal-*ϵ*), changed the somatic response. To break the symmetry between the two compartments in the distal–distal and proximal–proximal configurations, we added an external inhibitory input on both dendritic compartments. The strength of this input was proportional to the somatic activity (mimicking cortical feedback inhibition) and the input was injected by means of a conductance-based synapse, following Bono & Clopath (2017). The conductance was a double-exponential filter of the Tripod output spikes, with decay timescale of 50 ms, rise timescale of 2 ms, and peak conductance of 5 pS. The symmetry was broken by different strength feedback on the two dendrites. The B dendrite had a feedback peak-conductance that was *μ* times the baseline value, up to *μ* = 20, resulting in a peak conductance of 100 pS.

We tested whether the additional inhibitory feedback would make it possible to determine which of the two sequences was presented to the Tripod, AB*ϵ* or BA*ϵ*. We also compared the three dendritic configurations of the Tripod with a soma-only model. To distinguish the neural responses we calculated the average firing rate and the coefficient of variation of the inter-spike intervals (ISI CV) that can be used to detect burstiness. These spike statistics were computed during a period of 1 s, for 100 trials, in each of the four configurations. An example of the firing rate and ISI CV for a switch frequency of 6 Hz and *μ* = 5 is shown in Fig. 9*F* for the two sequences (orange and blue). We used logistic regression to quantify whether the two sequences could be distinguished. The outcome of a grid search over *μ* in the range 0 to 20 and switch frequencies of 1 Hz to 1000 Hz is shown in Fig. 9*G*. Classification accuracy was high for all dendritic configurations at switch frequencies below 100 Hz, and at chance level for the soma-only neuron. These findings were robust to variations in inhibitory feedback asymmetry (*μ*) and switch frequency. For the distal–distal model, large dendritic feedback inhibition reduced accuracy, likely because the Tripod did not spike enough to compute reliable statistics. For both symmetric configurations, accuracy was close to chance levels when feedback was the same on both dendrites (*μ* = 1). The proximal–distal model could recognize the sequences also when feedback was symmetric. This shows that sequence classification can be achieved reliably when neurons are equipped with dendritic compartments, whereas a single-compartment model (in its present instantiation) failed. Consistent with the previous results on switching, sequential order in the input could be distinguished based on the somatic spike response. The experiment used a fixed input rate and a fixed number of co-active synapses for both dendrites. It is to be expected that variability in rates and the number of input channels will increase the range of dendritic input patterns that can be decoded at the soma.

In conclusion, the Tripod model shows that neurons with dendrites have computational capabilities that single-compartment models lack. Cortical neurons, which receive thousands of spikes per second, can potentially use differences in the spatial location of the input to discriminate sequential information. Dendritic integration might be able to detect this variation and transfer the result of these local computations to the soma for downstream processing.

## Discussion

This paper has explored the computational implications of integrating dendritic compartments and voltage-gated receptors (NMDARs) into biological models of pyramidal neurons. We investigated the functional role of a simple dendritic structure in shaping the somatic response and analysed two classes of passive dendritic compartments, proximal and distal. The present work makes three main contributions. First, we have partitioned the space of dendritic morphology, connecting the emergence and dynamics of supra-linear integration to a small number of explainable geometric and physiological parameters. Secondly, our reduced neuron model performs dendritic computations that are usually reproduced only with more complex models.And third, we have outlined how dendrites contribute to structured computation, including logic operations, frequency detection, and sequence recognition. In summary, the relatively simple Tripod neuron proposes a reduced model of dendritic structure whose functionality transcends single-compartment models. The Julia implementation of the model can be readily used in lager-scale spiking neural network simulations.

In the first sections, we decomposed the model in minimal terms and investigated the contribution of various physiological and geometric factors in shaping the somatic and dendritic membrane dynamics. The comparison of human and mouse-like dendrites suggests that the former have longer integration timescales and are more excitable than their mouse counterparts, in agreement with experimental findings regarding the unique integrative properties of human dendrites (Fişek & Häusser, 2020; Beaulieu-Laroche *et al*., 2018; Beaulieu-Laroche *et al*., 2021). Our results confirm that human dendrites can be longer without losing the incoming current through membrane leakage; hence elongated geometries (distal thick) are possible, under our model’s constraints, with human but not with mouse parameters. The maximal length obtained for mouse is in agreement with basal and apical-oblique dendritic lengths in this species (Mohan *et al*., 2015). Later, we showed that independent of species-specific physiology, there is a geometric constraint that distinguishes between dendrites with strong agency on the soma (100 μm to 300 μm) and those with a slow and indirect action on it (300 μm to 500 μm). The theoretical distinction between distal and proximal dendrites in terms of the maximal elicited depolarization of the soma is consistent with previous experimental and computational work (Kamondi *et al*., 1998; Major *et al*., 2008; Eyal *et al*., 2018; Bono & Clopath, 2017) and it refers to the electronic distance between the dendritic compartment from the soma. Overall, dendritic lengths in the range of 100 μm to 500 μm correspond to dendrites in the basal and apical-oblique region of human pyramidal cells (Spruston, 2008). Passive dendrites and cable transmission are insufficient in modeling longer dendrites (e.g., apical-tuft of layer 2/3 and 5), suggesting that active, self-regenerative mechanisms such as calcium spikes (Larkum *et al*., 2007; Larkum, 2013) are required to transmit signals from distant dendritic input locations to the axon hillock. We associate the Tripod model to pyramidal cells rather than other types of cortical neurons for two main reasons. First, the physiological parameters adopted for both human and mouse cells and for both the membrane properties (Eyal *et al*., 2016; Dasika *et al*., 2007; Koch, 1999) and the NMDAR kinetics (Eyal *et al*., 2018; Duarte & Morrison, 2019) are obtained from electro-physiological studies on cortical pyramidal cells; While the interactions between dendritic integration and NMDAR non-linearity reported in the present paper could be valid for non-pyramidal cells, the different properties of NMDARs in spiny and non-spiny cells (Fleidervish *et al*., 2021; Augustinaite *et al*., 2014; Booker & Wyllie, 2021) may require ad-hoc model adjustments. Second, the dendritic lengths considered in the present work exceed those of other non-pyramidal cortical cells, such as layer IV spiny stellate cells (Meyer *et al*., 1989) and aspiny cells (Maxwell *et al*., 2007).

We also presented a detailed analysis of the somatic excitatory post-synaptic potentials (EPSPs) when inputs are received on distal and proximal dendrites and investigated synaptic efficacy and timescales with parameters obtained from human (Eyal *et al*., 2018) and mouse (Duarte & Morrison, 2019; Avermann *et al*., 2012) *in-vitro* experiments. Our results suggest that human-like voltage-dependent receptors (NMDARs) on distal dendrites affect dendritic integration. If dendritic compartments are sufficiently segregated electrically (distal), then co-activation of neighboring synapses produces NMDA spikes and, consequently, EPSPs with a supra-linear dependence on the number of synaptic inputs. These results are in agreement with *in-vitro* empirical findings (Polsky *et al*., 2004; Branco & Häusser, 2011; Eyal *et al*., 2018; Kumar *et al*., 2018; Bono & Clopath, 2017). Both electrophysiology and detailed computational models have shown that dendritic NMDA spikes can also be triggered in proximal synapses (Mel, 1992; Major *et al*., 2008). NMDA spikes in proximal dendrites result in larger somatic depolarization than distal ones. A few proximal NMDA spikes can drive the neuron to spike, while several distal NMDA spikes are required. Since the Tripod has only two compartments, the axial conductance to both proximal and distal synapses has to be larger than in multi-compartment models with several dendritic branches to impact the somatic membrane potential. With the present parameters, the axial conductance between the proximal and somatic compartment is large enough to trigger somatic spikes with a single depolarized proximal compartment. Therefore, our model accounts only for NMDA-induced plateau in distal dendrites because the proximal compartment can never reach the NMDA voltage-gating non-linerarity without triggering a bursty response in the soma; however, this does not result in a loss of generality for our model because the amplitude of somatic depolarization remains graded with respect to the dendritic length and it is weaker for longer dendrites, as measured experimentally *in-vitro* (Major *et al*., 2008). Because there are only two dendritic branches in the Tripod model, we have to interpret the axial currents, the NMDA-spikes, and the plateaus of dendritic Tripod’s compartments as an effective model of simultaneous depolarization in several dendritic branches of a pyramidal cell; crucially, recent evidence *in-vivo* has shown that the depolarization of a single hemi-tree of a pyramidal apical tuft, in contrast to both hemi-trees, have consequences in the behavioral scale Otor *et al*. (2022). In the current literature, there is considerable variability on the parameters used to replicate NMDA spikes, in particular in the choice of the NAR, which specifies the relative difference between the peak conductances of NMDA and AMPA receptors. For example, the NAR was set to 0.25 in Duarte & Morrison (2019), 1 in Bono & Clopath (2017), 1.2 in Ujfalussy & Makara (2020), 2 (Jadi *et al*., 2012), and 9 in Mel (1992). Empirical evidence, obtained mostly through indirect measurements, does report a similar level of variability. For example, NAR was found to be ≈ 0.25 and constant throughout the dendritic tree for mice hippocampal pyramidal neurons (Strube *et al*., 2017), roughly constant across different areas of the mouse neocortex (Myme *et al*., 2003), and NAR was ≈ 1.8 for human neocortical L2/3 pyramidal cells (Eyal *et al*., 2018). In this same spirit, we can interpret the discrepancy between the absence of NMDA spikes in mouse-like Tripod models and the experimental evidence of NMDA-related dendritic non-linearity in mice neurons Schiller *et al*. (2000); Antic *et al*. (2010); Larkum *et al*. (2022). Rather than postulating qualitative differences between mice and humans’ cells, we take it as an indication of minimal requirements for the emergence of NMDA-spikes in terms of timescales and steepness of NMDARs. In this respect, the variability in NMDA timescales of reduced models used in previous experiments dwarfed the difference in the NAR (50 ms (Bono & Clopath, 2017) 18.8 ms Jadi *et al*. (2012) 100 ms (Duarte & Morrison, 2019)). Our model identifies minimal geometric and NMDARs conditions for the occurrence of NMDA-spikes and emphasizes that merely implementing NMDA receptors is not sufficient for their emergence.

Previous computational models have found that somatic EPSPs are enhanced when inputs target different, independent dendrites (Dasika *et al*., 2007; Li *et al*., 2019), in an apparent conflict with experimental and computational evidence on synaptic clustering (Winnubst *et al*., 2015; Kastellakis *et al*., 2016; Bono & Clopath, 2017). This raises the question whether reduced models with passive dendritic compartments are sufficiently expressive to capture dendritic integration. Our results suggest that coincidence detection can be observed under certain conditions related to the location of synaptic input and synaptic physiology. The term coincidence-detection is used to characterize several dendritic phenomena (Spruston, 2008), e.g., the generation of a spike, or an activity burst, following simultaneous excitatory inputs. In particular, it is used for both the somatic depolarization resulting from simultaneous spikes on two segregated dendrites (Dasika *et al*., 2007), and for the non-linear response resulting from co-activation of neighboring synapses (Mel, 1992; Ujfalussy & Makara, 2020). Our model can express both forms of dendritic coincidence detection in terms of a single variable, i.e., dendritic length. The fundamental role of dendritic length has been discussed in Jadi *et al*. (2014) and was included in their two-layer network model of dendritic integration. However, the model only accounted for neuronal firing rates and did not model sub-threshold membrane dynamics. In addition, we explored the differences between inhibitory input onto the somatic and dendritic compartments. We associated dendritic inhibition with the activity of somatostatin interneurons (SST), and somatic inhibition to parvalbumin interneurons (PV) (Tremblay *et al*., 2016; Huang & Paul, 2019). From our fit on guinea pig pyramidal neurons (Miles *et al*., 1996), the GABA_A_receptors on the dendritic membrane had longer timescales and their maximal conductance was smaller than their somatic counterparts. We tested the differences between these two types of inhibition by comparing their efficacy in attenuating the somatic EPSP and showed that inhibition on the soma was effective in preventing spiking activity for a short period of time. In contrast, dendritic inhibition could silence the neuron for longer durations when applied on the same dendritic branch as excitation, but its maximal effect on the soma was limited and delayed, consistent with the current understanding of somatic and dendritic inhibition. The fast-spiking PV interneurons acting on the soma are associated with feed-forward, time-precise inhibition, while the slower action of SST cells regulates the dendritic potential via feedback inhibition (Tremblay *et al*., 2016; Kee *et al*., 2015; Tepper *et al*., 2008). In computational terms, localized inhibition allows for external gating of the dendritic stimulus by selecting which dendritic pathway is allowed to integrate the signal, and to communicate with the soma. Pathway-selection has been proposed as a cortical mechanism for flexible routing of sensory stimuli (Yang *et al*., 2016; Zajzon *et al*., 2019) and, more recently, it has been demonstrated that networks that leverage dendritic gating support efficient, durable, and fast learning (Sezener *et al*., 2021).

The Tripod succeeds in expressing coincidence-detection and pathway-selection because of two fundamental properties of its reduced dendritic tree: non-linear integration and electronic segregation of dendritic compartments. Our principled dendritic reduction aligns well with results from data-driven reductions that have been used to distill dendritic computations in the simplest architecture that could explain the data (Wybo *et al*., 2021; Beniaguev *et al*., 2021; Ujfalussy *et al*., 2018); in particular with the work by Ujfalussy *et al*. (2018) which shows how two compartments with non-linear integration and different timescales are sufficient to predict with high accuracy neural response under *in-vivo* stimuli conditions. However, dendritic simplification comes at a cost, synapses have no spatial resolution in the dendritic compartments but are all lumped together. Conversely, real dendrites are spatially extended and host spines, receptors, and ionic channels throughout the entirety of the dendritic cable. The interaction between synapses is determined by their relative distance and their spatial organization govern homeostatic mechanisms and heterosynaptic plasticity (Wu *et al*., 2020; Oh *et al*., 2015; Triesch *et al*., 2018; Kirchner & Gjorgjieva, 2021). The continuous spatial distribution along the dendritic cable also has important implications for signal integration: single-branch synaptic activation that follows the dromic direction - from the tip towards the soma - results in stronger somatic depolarization than activation in anti-dromic directions (Branco *et al*., 2010). Such distinctions are impossible under the constraints of our model, as we neglect spatial interactions along elongated dendrites comprising mutliple compartments. In addition, considering only two compartments bounds the computations available to each Tripod model to the dendritic configuration instantiated in the model, e.g. symmetrical or asymmetrical. In the brain each cell has hundreds of dendritic branches with a broad distribution of lengths, spatial arrangements and membrane physiology. Overall, the Tripod has to be considered as a compromise between accurate modelling of dendritic processes and implementing them in large-scale cortical circuits. As such, it provides a step forward from point-neuron models.

Dendritic NMDA spikes cause a long-lasting depolarization in the somatic compartment of the Tripod neuron. The duration of the depolarized state depends on dendritic length and the strength of synaptic events and it could last on the order of 100 ms, in agreement with experimental results (Schiller *et al*., 2000; Major *et al*., 2008; Milojkovic *et al*., 2005; Branco & Häusser, 2011). This dendritic “UP-state” is governed by a self-regenerative process triggered by co-active synapses and has a timescale that is two to three times longer than the membrane’s. This allows the UP-state to encode information about recent activity and the maintenance of this information can support an activity-silent processing memory at the neuronal level (Stokes, 2015; Fitz *et al*., 2020). Dendritic memory is similar to priming in the sense that the neuron responds faster and more strongly to a retrieval cue when the encoding signal occurs close in time. In contrast to short-lived synaptic memory (Mongillo *et al*., 2008), dendritic memory is more effective when the retrieval cue follows a different synaptic pathway than information encoding. The plateau potentials that support dendritic memory have been considered a candidate mechanism for linking neuronal to behavioral timescales (Augusto & Gambino, 2019; Bittner *et al*., 2015, 2017). Dendritic memory can bind information over time, and our results suggest that it can play a role in temporal processing that is beyond single-compartment models. To the question recently propose by Larkum (2022), *Are dendrites conceptually useful?* We can answer yes, because they naturally introduce a slow-decaying memory that facilitates the integration of sequential inputs in the timescale of 50 ms to 150 ms; which is extremely important for short-lived stimuli, such those in visual and auditory perception.

Since the introduction of the NEURON simulator in 1989 (Hines, 1989), the tools for modeling dendrites have come a long way (Poirazi & Papoutsi, 2020) and the incorporation of dendritic integration in cortical circuits is becoming increasingly accessible to computational research. Example of these advances are Dendrify (Pagkalos *et al*., 2022) and NESTML Plotnikov *et al*. (2016), which allow for simulating neurons with dendrites in cortical circuits in Brian2 and NEST. The Tripod model can be easily replicated within these frameworks. In addition, recent technical advances in neuromorphic computing have successfully implemented passive dendritic compartmentalization in hardware (Kaiser *et al*., 2022; Yang *et al*., 2021), boosting the applicability of dendritic computation in machine-learning contexts (Guerguiev *et al*., 2017; Sezener *et al*., 2021).The work presented here can guide this line of implementational research as it provides a simple, scalable model that captures important computational primitives at the single neuron level beyond the point neuron.

## Appendix A

### Minimal axial conductance

In order to simplify the analytical treatment, we consider the fixed point of the LIF equation, removing the exponential and the spike non-linearity. Because the slope of the AdEx nullcline is monotonous after *V*_*s*_ *> V*_*T*_, there is no qualitative difference in the presence of stationary input. Additionally, we consider a neuron driven solely by excitatory inputs. With two dendrites (*i* = 1, 2), the reduced tripod circuit is described by the following system of equations:

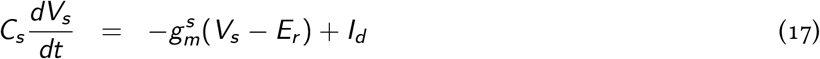

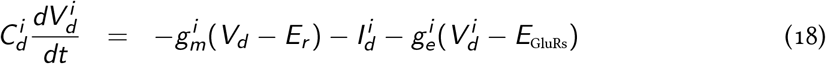

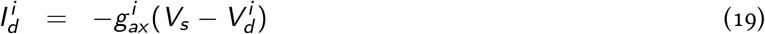

The system can be solved algebrically and, for *E*_GluRs_ = 0, results in:

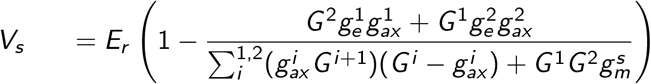

Where 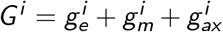. If the conductance between one of the dendrites and the soma is zero (neuron with single dendrite), the equation reduces to:

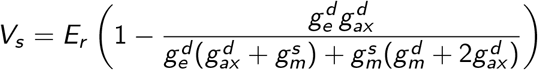

In the limit of very large excitatory conductances (*g*_*e*_ ≫ *g*_*m*_ + *g*_*ax*_), the neuron is a simple voltage divider and the somatic potential is given by:

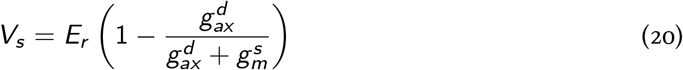

This situation corresponds to a neuron with a single dendrite and maximally excited in the

*d* -th dendritic compartment. Hence, the condition for the neuron to reach the spike threshold

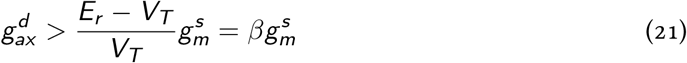

where *V*_*T*_ is the firing threshold of the somatic compartment. Eq. 13 defines the minimal condition for the dendritic compartment to elicit somatic spikes. When the axial conductance 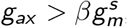, a full depolarization of the dendrites suffices to generate spikes in the soma.

Within the constraints of the Tripod model, some relevant parameters are fixed by the axosomatic model used, namely the somatic leak conductance 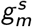, the resting membrane potential *E*_*r*_, and the spike threshold *V*_*T*_, which are all defined by the AdEx model Brette & Gerstner (2005). The remaining parameter for the axial conductance 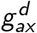 is determined entirely by the cable geometry (diameter *d* and length *l*) (Rall, 2011) along with the dendritic membrane physiology as expressed in Eqs. 5 6, 7. Once the physiological details are defined (Dasika *et al*., 2007; Eyal *et al*., 2016), we can distinguish between geometries that elicit spikes and geometries that do not.

## Appendix B

### Excitatory synaptic interactions in the passive cable

Dasika *et al*. (2007) shows that a model neuron with stationary conductance depolarizes more when the inputs are distributed than when synapses are localized on a single branch. This can be demonstrated by determining the equivalence between a circuit with two active synapses on different branches (*G*_1_ and *G*_2_) and a circuit with one single active conductance *G*_*s*_ (*G*_1_ = *G*_*s*_ and *G*_2_ = 0). The following equivalence holds:

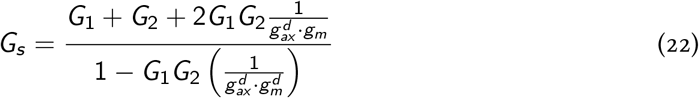

where *g*_*ax*_ and *g*_*m*_ are the axial and the leak conductances of the passive membrane patch (the dendritic compartments), respectively. The equation shows that, in the presence of segregated dendritic compartments (*g*_*ax*_ *<* ∞), *G*_*s*_ is a always greater than *G*_1_ + *G*_2_. The interaction has been further simplified in (Li *et al*., 2019). The authors reduce the interaction between synapses in a second order approximation where the total current, in case of simultaneous firing synapses, is given by:

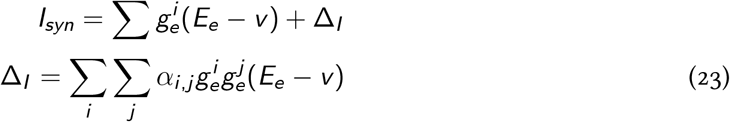

where the *E*_*e*_ states the receptor reverse potential. The interaction has been approximated to a binary function where the second order synaptic contribution *α*_*i,j*_ is almost zero for synapses on different branches and negative for synapses on the same branch.

## Appendix C

### Membrane dynamics across experiments

#### Balanced inputs condition

To study the model in naturalistic conditions we stimulated the Tripod with excitatory and inhibitory spike trains. We defined a balanced condition such that the somatic compartment is depolarized and both glutamatergic and gabaergic conductances are large; this input configuration ensures that the dendritic computations investigated are not artifacts of the unrealistic set-up. The balance is obtained by fixing the excitatory firing rate to 3 kHz and varying the corresponding inhibitory rates. This procedure results in inhibitory firing rates of 3 kHz for distal dendrites (400 μm); 4.8 kHz for proximal dendrites (150 μm); and 1 kHz for the soma only model. With these inputs, the neuron (almost) never fires and the somatic compartment rests around −67 mV for the three dendritic configurations, distal-distal, distal-proximal, and proximal-proximal. Following the protocols presented in the Results section, each dendrite was activated by doubling the excitatory input; when this happens, the dendrite depolarizes and causes the neuron to fire. When both dendrites are activated the neuron’s firing rate is approximately 30 Hz, with little variations between different dendritic configurations. In experiments Fig. 7, Fig. 8, Fig. 9 additional excitatory noise was injected in the somatic compartment to ensure firing activity when one single dendrite was activated.

The *soma only* balanced configuration was also defined on a similar basis, although the soma compartment needs to be more depolarized −60 mV to initiate spikes when one of the input pathways is activated. Fig. 10 illustrates these effects and shows the three Tripod configurations and the soma-only condition in the inactive (A) and active (B) states.

**Figure 10:**
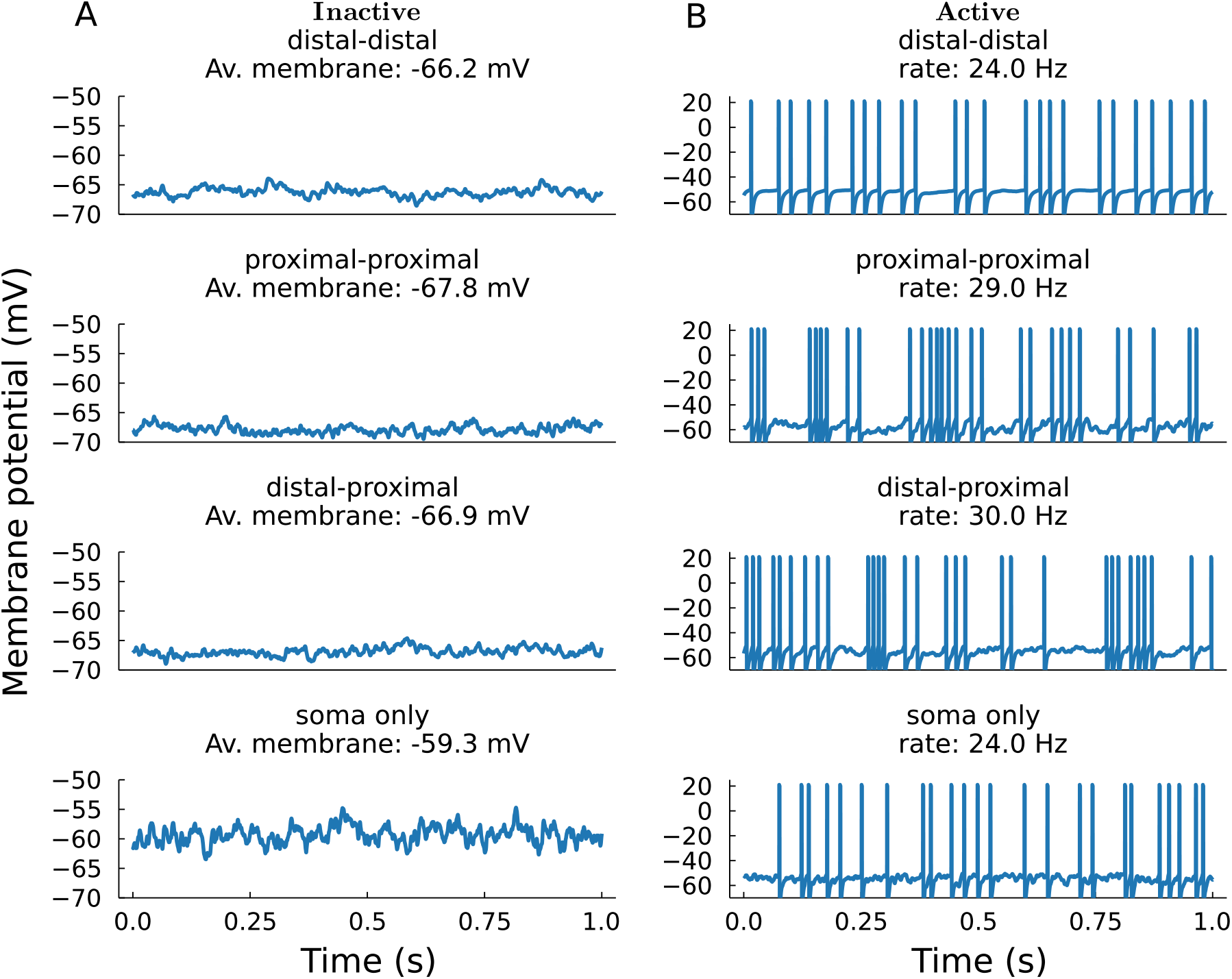
Membrane dynamics of Tripod models. (**A**) Model activity in the inactive condition, with 3 kHz excitatory inputs and dendritic-length-specific values for inhibition (see main text). (**B**) Membrane dynamics in the active mode with doubled excitatory input rates.

An important outcome of the balanced configuration is to avoid artifacts of the AdEx model, as discussed in Górski *et al*. (2021). When the AdEx is strongly excited, for example with strong GluRs stimulation or injected currents, the neuron starts firing and the adaptive current rapidly rises. If the stimulation terminates abruptly, the adaptive current pulls down the membrane voltage, generating unnatural hyper-polarization. In Fig. 11,Fig. 12,Fig. 13 we show that, due to our balance condition, these artifacts are not observed in the Tripod model and realistic membrane dynamics can be observed.

**Figure 11:**
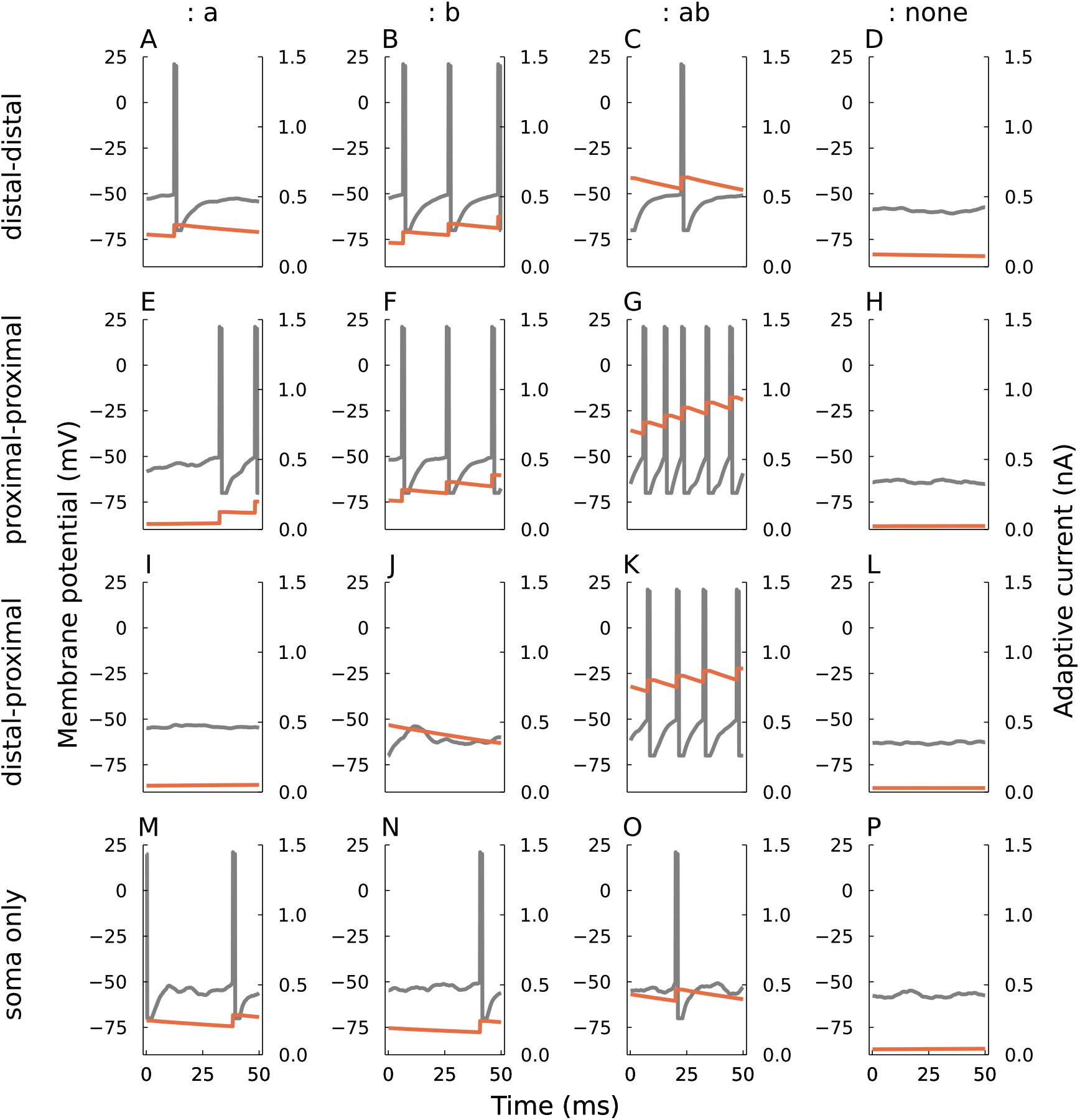
Tripod dynamics in the Logical operators task. **(A)**-**(P)** Somatic membrane potential (gray) and adaptive current (red) for the 4 dendritic configurations (vertical arrangement) and the 4 input configurations (horizontal arrangement).

**Figure 12:**
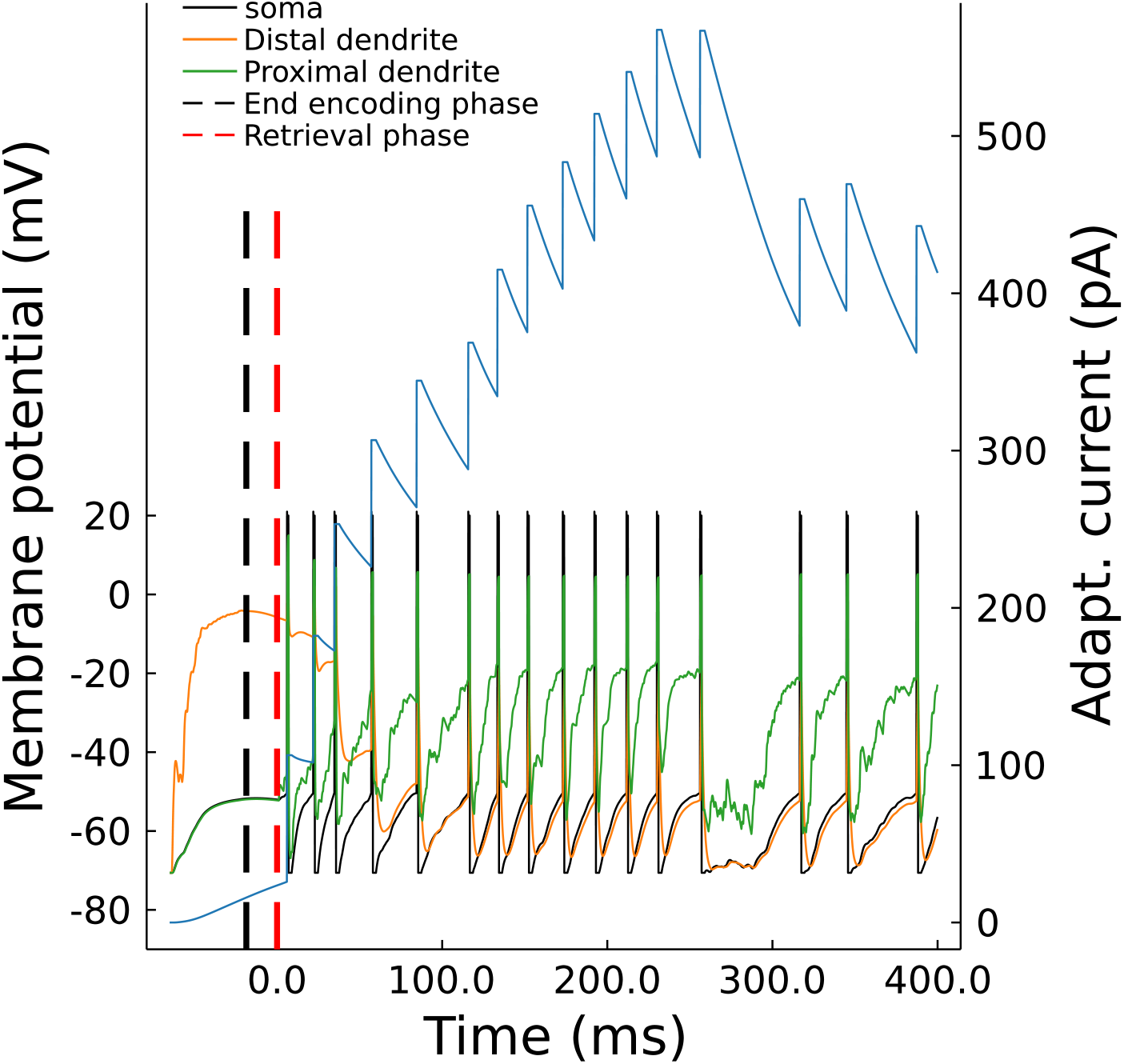
Tripod dynamics in the Dendritic memory operators task. The figure depicts somatic membrane voltage and adaptive current (black and blue) as well as dendritic voltages (red and green) for the distal-proximal dendritic configuration. Vertical dashed lines in red and black indicate the end of the encoding phase and the onset of the retrieval phase, see main text for clarifications.

**Figure 13:**
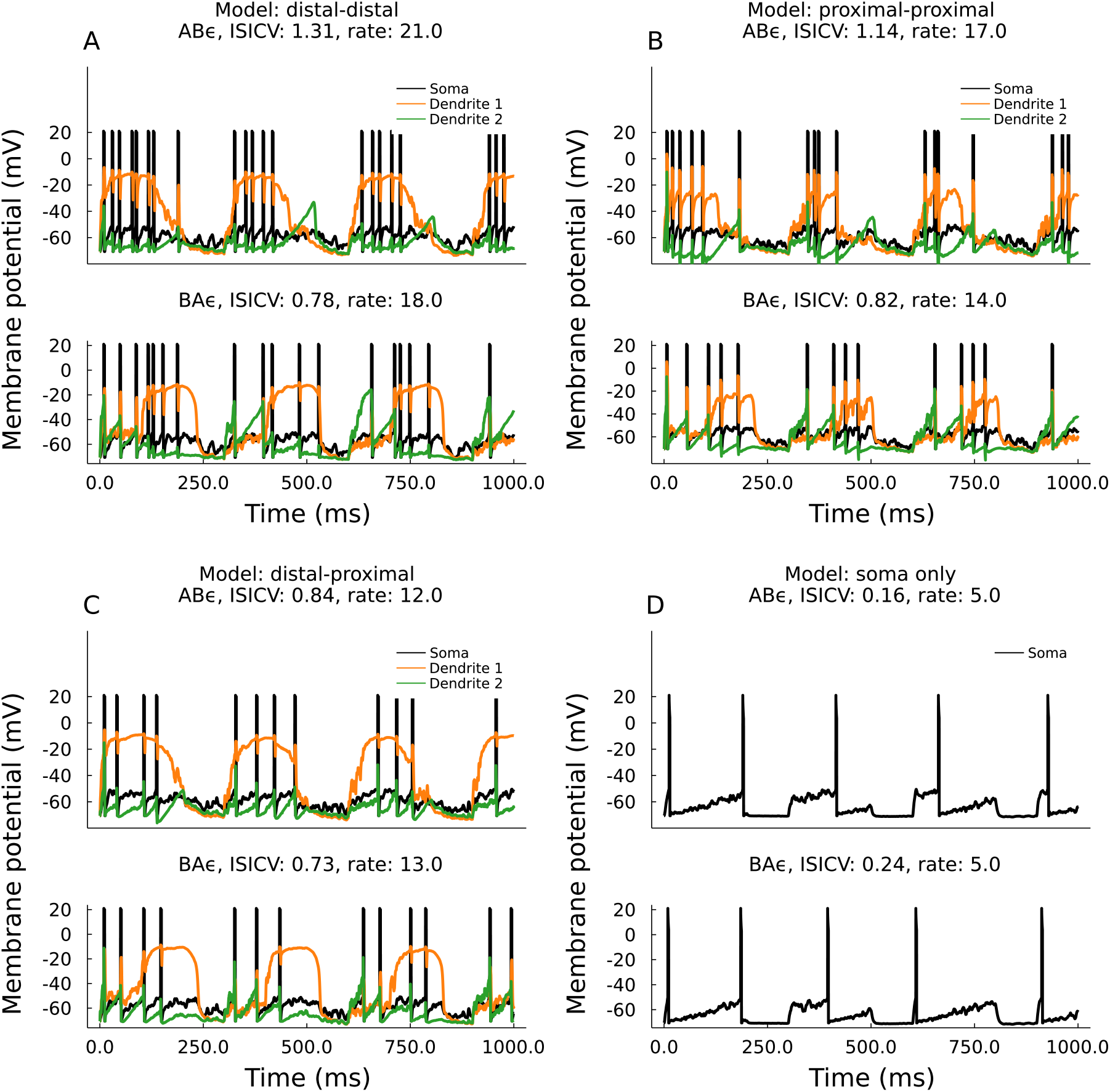
Tripod dynamics in the Sequence recognition task. Somatic membrane potential (black) and dendritic membrane potentials (red and green) for the four Tripod configurations. The two panels in **(A)** - **(D)** show the dynamics of the Tripod in the AB*ϵ* and the BA*ϵ* sequence, see main text. To facilitate the comparise, these simulations were run with frozen noise input. As such, the difference between the upper and lower panels is only the strength of the excitatory inputs which is doubled in the stimulated compartment: input A in the AB sequence (top pannels) and input B the BA sequence (bottom pannels).

## Notes

### Competing Interest Statement

The authors have declared no competing interest.

